# Erythropoietin-dependent Acquisition of CD71^hi^CD105^hi^ Phenotype within CD235a^−^ Early Erythroid Progenitors

**DOI:** 10.1101/2024.08.29.610192

**Authors:** Natascha Schippel, Jing Wei, Xiaokuang Ma, Mrinalini Kala, Shenfeng Qiu, Peter Stoilov, Shalini Sharma

## Abstract

The development of committed erythroid progenitors and their continued maturation into mature erythrocytes requires the cytokine erythropoietin (Epo). Here, we describe the immunophenotypic identification of a unique Epo-dependent colony-forming unit-erythroid (CFU-E) cell subtype that forms during early erythropoiesis (EE). This previously undescribed CFU-E subtype, termed late-CFU-E (lateC), lacks surface expression of the characteristic erythroid marker CD235a (glycophorin A) but has high levels of CD71 and CD105. LateCs could be prospectively detected in human bone marrow (BM) cells and, upon isolation and reculture, exhibited the potential to form CFU-E colonies in medium containing only Epo (no other cytokines) and continued differentiation along the erythroid trajectory. Analysis of *ex vivo* cultures of BM CD34^+^ cells showed that acquisition of the CD7^hi^CD105^hi^ phenotype in lateCs is gradual and occurs through the formation of four EE cell subtypes. Of these, two are CD34^+^ burst-forming unit-erythroid (BFU-E) cells, distinguishable as CD7^lo^CD105^lo^ early BFU-E and CD7^hi^CD105^lo^ late BFU-E, and two are CD34^−^ CFU-Es, also distinguishable as CD71^lo^CD105^lo^ early CFU-E and CD7^hi^CD105^lo^ mid-CFU-E. The transition of these EE populations is accompanied by a rise in CD36 expression, such that all lateCs are CD36^+^. Single cell RNA-sequencing analysis confirmed Epo-dependent formation of a CFU-E cluster that exhibits high coexpression of CD71, CD105, and CD36 transcripts. Gene set enrichment analysis revealed the involvement of genes specific to fatty acid and cholesterol metabolism in lateC formation. Overall, in addition to identifying a key Epo-dependent EE cell stage, this study provides a framework for investigation into mechanisms underlying other erythropoiesis-stimulating agents.

## Introduction

Differentiation of hematopoietic stem cells (HSCs) to erythrocytes or red blood cells (RBCs) is dependent on erythropoietin (Epo), a cytokine produced and released by the kidneys^1-3^. Epo is necessary for the survival and continued differentiation of erythroid-lineage-committed progenitor colony-forming unit-erythroid (CFU-E) cells in both humans and mice^4,5^. Mice lacking Epo or its receptor (EpoR) die by embryonic day 13.5 due to a lack of mature RBCs^6-9^. Binding of Epo induces homodimerization of EpoR, which stimulates the receptor-associated Janus Kinase 2 (JAK2) to phosphorylate EpoR and itself^10,11^. This process is associated with downstream activation of three hallmark signal transduction pathways: signal transducer and activator of transcription 5 (STAT5), mitogen-activated protein kinase/extracellular signal-regulated kinase (MAPK/ERK), and phosphoinositide-3 kinase/protein kinase B (PI3K/Akt)^2,12-15^. Activation of these pathways induces the expression of genes that inhibit apoptosis and promote survival, proliferation, and continual differentiation of CFU-Es.

Classically, the development of erythrocytes from HSCs has been described as a multistage process that progresses through intermediate progenitors, including multipotent progenitors (MPPs) and common myeloid progenitors (CMPs)^16-20^. The latter give rise to the bipotent megakaryocyte-erythroid progenitors (MEPs), from which unipotent erythroid progenitor burst-forming unit-erythroid (BFU-E) cells arise. The formation of BFU-E from MEP and their conversion to CFU-E, a phase referred to as early erythropoiesis (EE), is followed by terminal erythroid differentiation (TED). The transition of CFU-Es to proerythroblasts (ProEs) marks the onset of TED, which advances along ProEs, basophilic erythroblasts (BasoEs), polychromatic erythroblasts (PolyEs), and finally, orthochromatic erythroblasts (OrthoEs). The enucleation of OrthoEs forms reticulocytes, which ultimately mature into RBCs. Early single cell RNA-sequencing (scRNA-seq) analyses indicated a continuum of differentiation from HSCs to unipotent progenitors and the emergence of unipotent progenitors of erythroid and other lineages directly from multipotent stem cells rather than a stepwise hierarchical process^21-23^.

Recent computational analysis of an extensive dataset comprising scRNA-seq data for ∼260,000 bone marrow (BM) cells from six different studies has revealed the existence of distinct transcriptional states corresponding to clusters of multipotent (LMPP, MPP, and CMP), bipotent (GMP and MEP), and unipotent (BFU-E and CFU-E) progenitors, as well as EE and TED populations along the erythroid differentiation pathway^24^. Similar transcriptional profiling of healthy young and elderly individuals has identified multi- and bipotent progenitor cell-specific clusters along the differentiation trajectory^25^. Longitudinal single cell proteomic analysis has also detected all erythroid developmental stages but suggests lineage choice to be a gradual process rather than a quick fate-determining decision^26^. Measurement of protein levels showed that transcription factors from two lineages may be coexpressed in bipotent progenitors, and a slow quantitative increase in lineage-specific transcription factors underlies commitment.

Based on a core set of surface markers, several immunophenotyping strategies have been proposed for the analysis of erythroid stages^27-35^. These strategies employ markers that are expressed during the differentiation of other lineages, although the critical role of many in erythroid differentiation is well established. They have been especially useful in monitoring transitions during EE and TED, and include early markers that are present in hematopoietic stem and progenitor cells (HSPCs) and continue to be expressed into EE or TED, such as CD117 (c-kit), and CD49d (α-4 integrin). Other markers are transient, such as CD36 (fatty acid translocase), CD105 (endoglin), and CD71 (transferrin receptor), or arise late and are maintained in mature RBCs, such as CD235a (glycophorin A) and CD233 (Band 3). Conventionally, the BFU-E to CFU-E transition has been associated with loss of CD34 expression, gain of CD36 expression, and increase of CD71 and CD105 expression^28,29,31,36^. While some studies have reported expression of CD71 and CD105 in bipotent MEPs, others have suggested that CD71 and CD105 arise later, at or just before commitment to the erythroid lineage in EE cells^29,31-33^. Finally, immunophenotypic analysis has revealed heterogeneity within EE populations^28^. Based on CD34 and CD105 expression on CD235a^−^CD117^+^CD45RA^−^CD41a^−^CD123^−^CD71^+^ cells, Yan et al. resolved four distinct subsets of erythroid progenitors, including one BFU-E (EP1) and three CFU-E populations (EP2-4). Proteomic analysis has also detected four CFU-E populations, two of which were CD34^−^CD45RA^−^CD123^−^CD36^+^CD71^+^CD235a^−^ but differed based on CD38 positivity; CFU-E1 were CD38^+^ and CFU-E2 were CD38^−26^. The other two, CFU-E3 and CFU-E-like, expressed the same combination of markers as CFU-E2, but at a lower level, and the CFU-E-like lacked CD36 and CD71. However, exactly which of these subpopulations is Epo-dependent and how Epo regulates the dynamics of transition within these EE subtypes has not yet been investigated.

In this study, we utilized a combination of immunophenotyping and scRNA-seq to assess the influence of Epo on EE population dynamics during *ex vivo* differentiation of human BM CD34^+^ HSPCs. Based on profiles of established erythroid surface markers, we developed an immunophenotyping strategy for the delineation of BFU-E and CFU-E cells. Application of this strategy to HSPCs undergoing differentiation revealed five EE populations, which could also be resolved prospectively in BM cells: early BFU-E (earlyB), late BFU-E (lateB), early CFU-E (earlyC), mid CFU-E (midC), and late CFU-E (lateC). Additionally, comparison of cells undergoing differentiation in the presence and absence of Epo identified the acquisition of the CD71^hi^CD105^hi^ immunophenotype during the conversion of midC to lateC as the crucial Epo-dependent step. Our strategy also enabled the isolation of the EE subtypes by fluorescence activated cell sorting (FACS) and their further characterization by reculturing and colony forming cell (CFC) assays to confirm their ability to continue differentiation along the erythroid trajectory. Analysis by scRNA-seq validated the Epo-dependent transition to high coexpression of *TFRC* (CD71) and *ENG* (CD105) transcripts and progression to TED. Transcriptomic profiles of the differentially expressed genes (DEGs) detected the established erythroid differentiation genes and pathways—such as those involved with heme and globin synthesis, iron uptake, and the erythroid transcription factors—and identified several new Epo-induced genes, including those involved in lipid metabolism. Overall, this study provides a framework for the assessment of EE populations and transcriptomic alterations that can be applied to investigating the influence of factors on the regulation of erythropoiesis.

## Results

### Epo-associated Dynamics of CD71 and CD105 Expression

To examine Epo-associated cellular dynamics during erythroid differentiation, we applied a previously established protocol for *ex vivo* differentiation of human BM-derived CD34^+^ HSPCs and performed longitudinal immunophenotypic analysis to assess differentiation^37-39^. As CD235a is known to first arise at the onset of TED, we assessed CD235a expression in cells cultured in medium containing Epo (+Epo) or lacking Epo (−Epo). While >50% of live cells were CD235a^+^in +Epo cultures, there was a lack of a significant CD235a^+^ population in −Epo medium (Figures S1A and S1B). Additionally, assessment of erythroblast populations using the flow cytometry strategy developed by Yan et al. showed the presence of all five erythroblast populations (ProE, early BasoE, late BasoE, PolyE, and OrthoE) in +Epo but no erythroblasts undergoing TED in −Epo cultures (Figure S1C)^28^.

Next, we examined the expression of markers that have previously been used to distinguish and characterize EE and TED populations, including CD49d, CD117, CD36, CD71, and CD105^16,28-31,33,40^. We found that the percentage of LiveLin^−^ cells expressing CD105 (days 4 and 7), CD71 (day 4), and CD36 (days 4, 7, and 10) was significantly reduced in −Epo cultures, while those expressing CD49d and CD117 were unaffected (Figures 1A, 1D, and S1D-S1I). Further, both CD105 and CD71 exhibited expression ranges encompassing low and high levels during the course of differentiation. This phenomenon was observed with CD105 and CD71 antibodies labeled with two different fluorophores and has been reported previously by others (Figures 1B, 1E, and S3A and S3B)^27,28,30-32,35,41^. Furthermore, the fraction of CD71^hi^ and CD105^hi^ cells was significantly reduced in −Epo cultures (Figures 1C and 1F). This decrease was accompanied by an increase in CD71^lo^ but not CD105^lo^ cells. Overall, the results confirmed the Epo-induced development of erythroblasts in our culture system and indicated the ability of Epo to influence the surface expression of CD71 and CD105.

**Figure 1:**
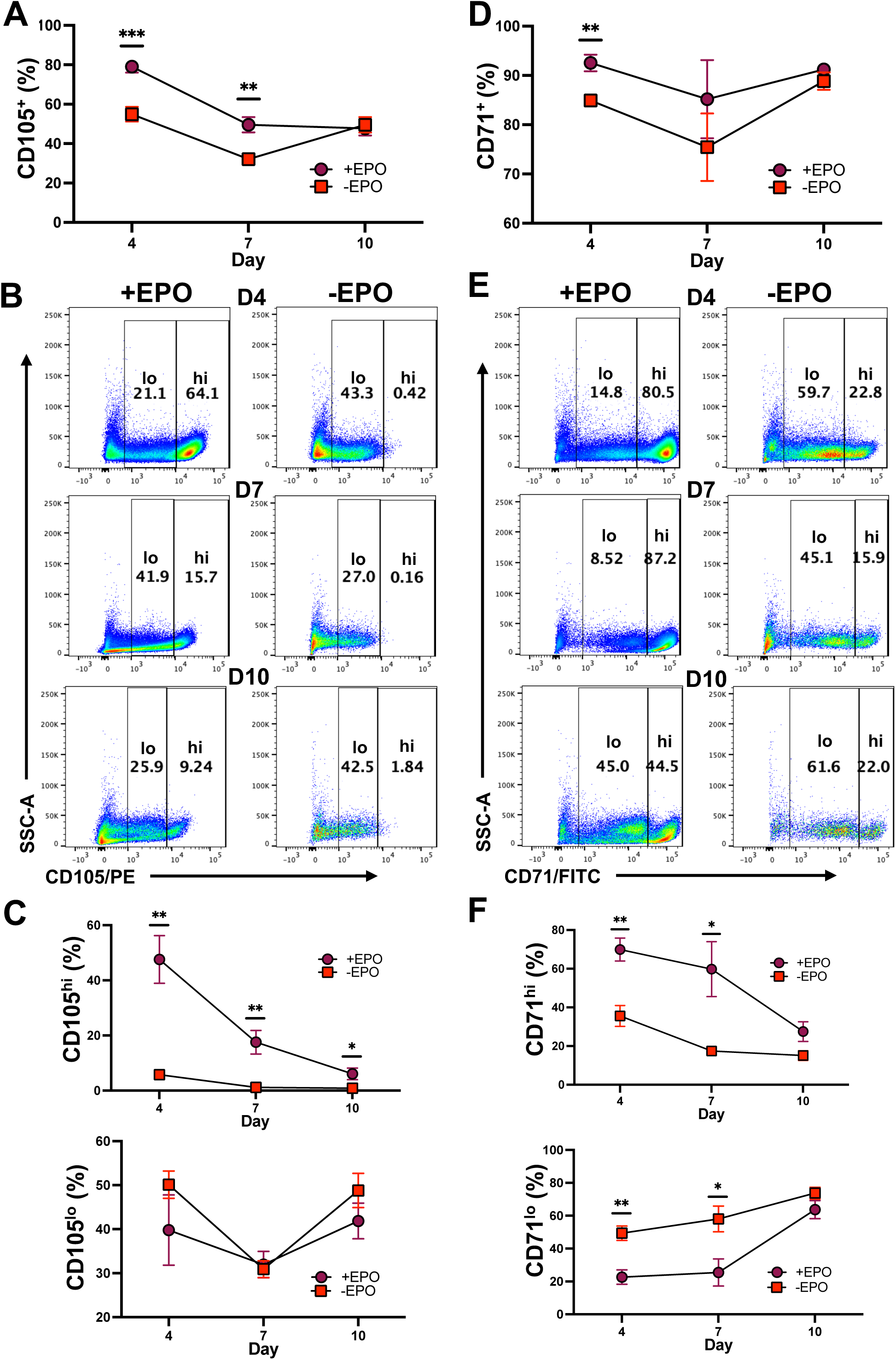
Epo induces high expression of CD105 and CD71. BM CD34^+^ HSPCs were cocultured with MS-5 stromal cells in medium containing or lacking Epo. Surface expression of CD105 and CD71 was determined by flow cytometry, and percentages of positive cells within live cells are shown. **(A)** Overall expression of CD105 on cells from BM HSPC cultures with and without Epo. **(B)** Representative scatter plots for CD105 expression in +/− Epo cultures, indicating gates for low and high expression as determined by anti-CD105/PE antibody staining. **(C)** Comparison of percentages of CD105^hi^ (top) and CD105^lo^ (bottom) cells in +/−Epo cultures (n = 5). **(D)** Overall expression of CD71 on cells from BM HSPC cultures with and without Epo. **(E)** Representative plots for CD71 expression in +/− Epo cultures, indicating gates for low and high expression as determined by anti-CD71/FITC antibody staining. **(F)** Comparison of percentages of CD71^hi^ (top) and CD71^lo^ (bottom) cells in +/− Epo cultures (n = 5). P-values were calculated by the Student T-test, and values ≤ 0.05 were considered significant (*p < 0.05, **p < 0.01, ***p < 0.001). Abbreviations: Epo; erythropoietin.

### Immunophenotyping Strategy for Resolution of Early Erythroid Progenitors

Utilizing previously described markers, we developed an immunophenotyping strategy for monitoring the Epo-dependent transition of EE progenitors, BFU-Es and CFU-Es (Figure 2). For this, surface markers for CD3, CD10, CD19, CD11b, CD41a, and CD123 are used to gate out non-erythroid lineages and progenitors; CMP and GMPs are both expected to be CD123 positive and therefore should be eliminated with this panel. From the LiveLin^−^CD123^−^cells, CD71^+^CD235a^−^ immature cells were first selected and then analyzed for expression of CD49d and CD117, as these markers have been shown to be expressed in EE progenitors (Figure 2A i and ii)^28,31^. From this LiveLin^−^CD123^−^CD71^+^CD235a^−^ population, only CD49d^+^CD117^+^ double positive cells were considered for further analysis (Figure 2A ii). As BFU-Es, but not CFU-Es, are expected to express CD34, all CD105^+^ cells were segregated based on CD34 positivity to separate BFU-Es from CFU-Es (Figure 2A iii). Therefore, all EE cells were LiveLin^−^CD123^−^CD235a^−^CD49d^+^CD117^+^, and within this population, total BFU-Es were additionally CD34^+^, whereas total CFU-Es were CD34^−^. Further stratification of total BFU-E and CFU-E populations based on CD105 and CD71 expression resolved heterogeneity within them (Figure 2A iv and v). BFU-Es consisted of two populations: a large CD71^lo^CD105^lo^ population and another smaller population that was CD71^hi^CD105^lo^. Interestingly, within CFU-Es, three populations emerged: CD71^lo^CD105^lo^, CD71^hi^CD105^lo^, and CD71^hi^CD105^hi^. Thus, both BFU-E and CFU-E cohorts contained CD71^lo^CD105^lo^ and CD71^hi^CD105^lo^ populations that only differed by the expression of CD34, whereas a CD71^hi^CD105^hi^ cohort was only observed within the CD34^−^ CFU-Es.

**Figure 2:**
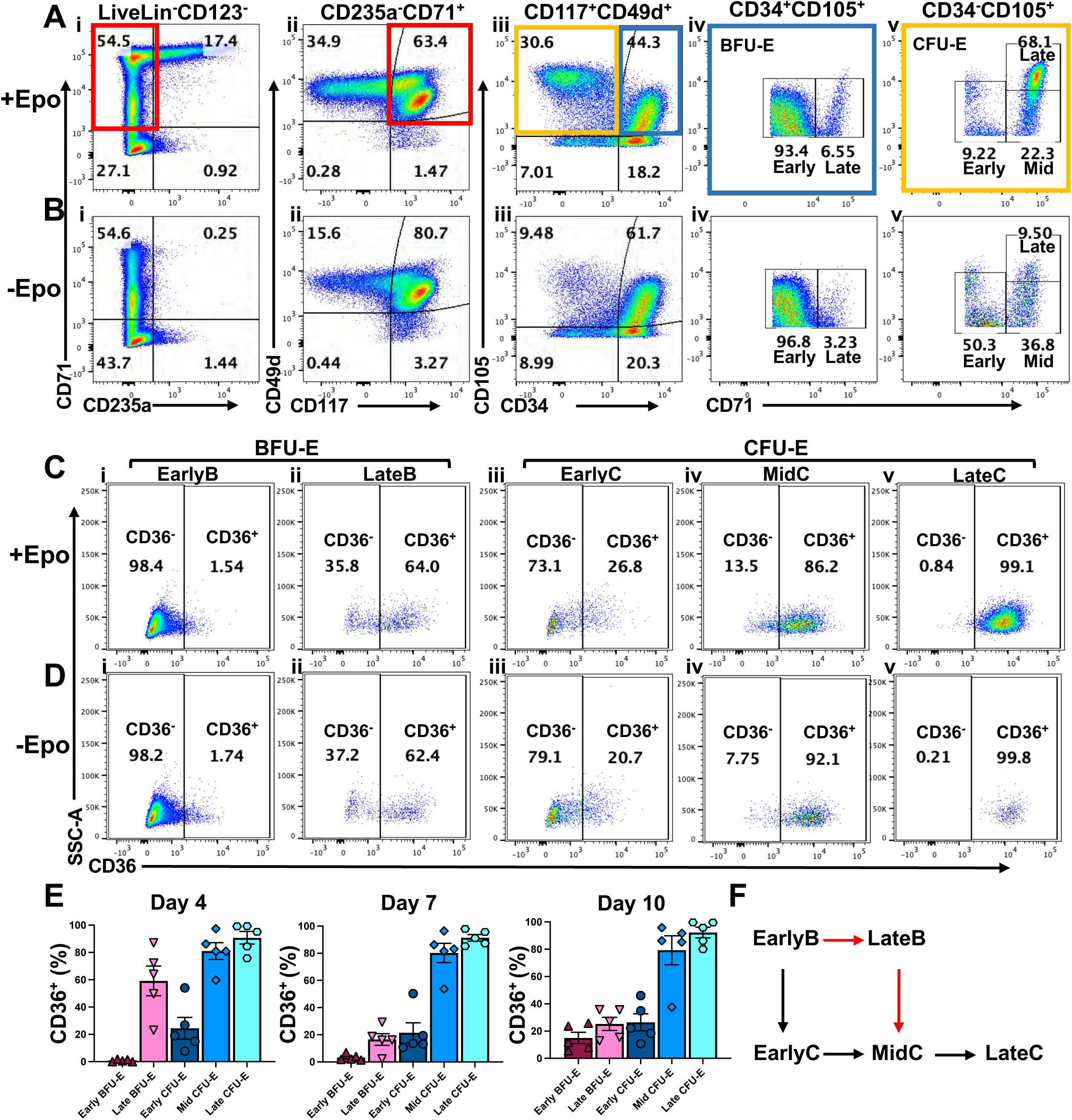
Immunophenotyping strategy for stratification of early erythroid progenitor populations. Cells from ex vivo cultures of BM CD34^+^ HSPCs incubated in medium containing or lacking Epo were analyzed for the indicated surface markers by flow cytometry. **(A)** Representative scatter plots from the analysis of heterogeneous EE populations in cultures containing Epo. CD235a^+^CD71^+^ cells are gated from LiveLin^−^ cells (i), then CD117^+^CD49d^+^ cells are selected (ii), and CD105^+^ cells are stratified based on CD34 expression (iii). All CD34^+^CD105^+^ cells are considered BFU-Es (iv), and all CD34^-^CD105^+^ cells are considered CFU-Es (v). Further stratification of BFU-E and CFU-E based on CD71 and CD105 expression showed two populations within BFU-Es: CD71^lo^CD105^lo^ (earlyB) and CD71^hi^CD105^lo^ (lateB) (iv). Within CFU-Es, three populations were observed: CD71^lo^CD105^lo^ (earlyC), CD71^hi^CD105^lo^ (midC), and CD71^hi^CD105^hi^ (lateC) (v). **(B)** Representative scatter plots for analysis of EE populations in cultures lacking Epo by the same strategy as in (A). **(C)** Representative scatter plots depicting CD36 expression within each of the five EE populations from cultures grown in medium containing Epo: i) EarlyB, ii) LateB, iii) EarlyC, iv) MidC, and v) LateC. **(D)** Representative scatter plots depicting CD36 expression within each of the five EE populations from cultures grown in medium lacking Epo: i) EarlyB, ii) LateB, iii) EarlyC, iv) MidC, and v) LateC. **(E)** Quantification of CD36 expression in all five EE populations on days 4, 7, and 10 (n = 5). **(F)** Proposed differentiation routes for the transition of progenitors in EE. All scatter plots are from flow cytometry analysis performed on day 4 of *ex vivo* culture. Abbreviations: Epo; erythropoietin, BFU-E; burst forming unit-erythroid, CFU-E; colony forming unit-erythroid, earlyB; early BFU-E, lateB; late BFU-E, earlyC; early CFU-E, midC; mid CFU-E, lateC; late CFU-E.

To further characterize the EE populations, we assessed CD36, which has previously been employed to distinguish BFU-E and CFU-E cells^29,42,43^ (Figure 2C). Amongst BFU-Es, the fraction of CD36^+^ cells was markedly higher in the CD71^hi^CD105^lo^ cohort (∼59%) than in the CD71^lo^CD105^lo^ cohort (∼1.5%) (Figure 2C i and ii, and 2E). While within CFU-Es, the percentage of CD36^+^ cells increased from ∼25% in CD71^lo^CD105^lo^ cells to ∼81% in CD71^hi^CD105^lo^ and ∼99% in CD71^hi^CD105^hi^ cells (Figure 2C iii-v, and 2E). Thus, based on the expression of CD34, CD71, CD105, and CD36, we assigned the terms early-BFU-E (earlyB) to the population that was CD34^+^CD71^lo^CD105^lo^ and had very few CD36^+^ cells, and late-BFU-E (lateB) to CD34^+^CD71^hi^CD105^lo^ cells that had a comparatively higher percentage of CD36^+^ cells (Figure 2C i and ii, and 2E). Similarly, the CFU-E populations were termed early, mid, and late: early-CFU-E (earlyC) were CD34^−^CD71^lo^CD105^lo^ with a low percentage of CD36^+^ cells and mid-CFU-E (midC) were CD34^−^CD71^hi^CD105^lo^ with an intermediate percentage of CD36^+^ cells, whereas late-CFU-E (lateC) were CD34^−^CD71^hi^CD105^hi^ and all CD36^+^ (Figure 2C iii-v, and 2E). These results indicate that within the total EE population, there is a continuum of cells that are simultaneously undergoing loss of CD34 and gain of CD36 expression. The observed patterns suggest the existence of two alternative differentiation routes (Figure 2F). In one route, loss of CD34 and gradual gain of CD36 transition earlyBs to earlyCs, that form midCs and lateCs. In the other route, the onset of CD36 expression precedes the downregulation of CD34, leading to the transition of earlyBs to lateBs. In this alternative route, the earlyC stage is likely bypassed, with lateB first forming midC and then lateC.

To confirm that the observed stratification of EE cells was valid and not an artifact of the combination of fluorophores that were used in our panel of antibodies, we designed a second panel in which CD49d, CD71, CD105, and CD36 antibodies conjugated to different fluorophores were used (Figure S3C and S3E). The list of antibodies for the two panels are provided in Table S1. Application of the second panel also demonstrated heterogeneity in EE cells, identifying the presence of five populations at percentages similar to those observed with the initial fluorophore combination (Figure S3C and S3G-3M). Additionally, the pattern of CD36 expression was also similar to that seen with the first panel (Figure S3E). Put together, these results demonstrate that heterogeneity within EE cells can be resolved based on CD71 and CD105 expression and that the rates of CD34 downregulation and CD36 upregulation may vary within each cell.

### Erythropoietin is Required for the Development of Late CFU-Es

To examine the role of Epo in population dynamics within EE, cells from BM-derived CD34^+^ HSPCs cultured in +Epo medium were compared to those cultured in −Epo medium by immunophenotyping. Assessment of total BFU-Es during the course of +Epo culture showed an increase from day 4 to 7, which was followed by maintenance of their levels until day 10; this trend was largely maintained in the absence of Epo (Figures 2A iii, 2B iii, and 3A). In line with this observation, percentages of earlyBs and lateBs were also unaffected by the absence of Epo (Figures 2A iv, 2B iv, 3B, and 3C). The levels of total CFU-Es decreased steadily from day 4 to 10 in +Epo conditions (Figure 3D). Although there was some variability in the trend exhibited by the individual erythroblast populations, CD235a expression indicated an increase in total erythroblasts, and the percentage of OrthoE, which is the final erythroblast stage, also increased steadily, thereby indicating a continuous transition into TED (Figures 3E, and 3F, and S1A-S1C). In the absence of Epo, however, there was a significant decrease in total CFU-Es on days 4 and 7 of culture (Figures 2A iii, 2B iii, and 3D). The lack of difference in CFU-E percentages in +Epo and −Epo conditions on day 10 is likely due to their progression into TED in the presence of Epo. Interestingly, examination of the individual CFU-E populations revealed differential effects within the subtypes. The percentage of lateCs was significantly lower in −Epo compared to +Epo cultures on all three days of assessment (Figures 2A v, 2B v, and 3I). Whereas the percentages of both earlyCs and midCs were higher in the absence of Epo than in the presence of Epo (Figures 2A v, 2B v, 3G, and 3H). These differences were significant only on day 4 and not at later timepoints, most likely due to the progression of CFU-Es to TED in +Epo cultures. These results suggest that a lack of Epo may be causing reduced production of lateCs with concomitant accumulation of earlyCs and midCs. In other words, the transition from midC to lateC and the acquisition of the CD71^hi^CD105^hi^ immunophenotype are dependent on the presence of Epo. The absence of Epo did not have a significant effect on BFU-Es or their transition to earlyCs or midCs. Analysis using the second antibody panel indicated similar percentages of EE cells and a lack of effect on BFU-Es but showed accumulation of earlyCs and reduced production of lateCs (Figures S3C-S3D, and S3G-S3M).

**Figure 3.**
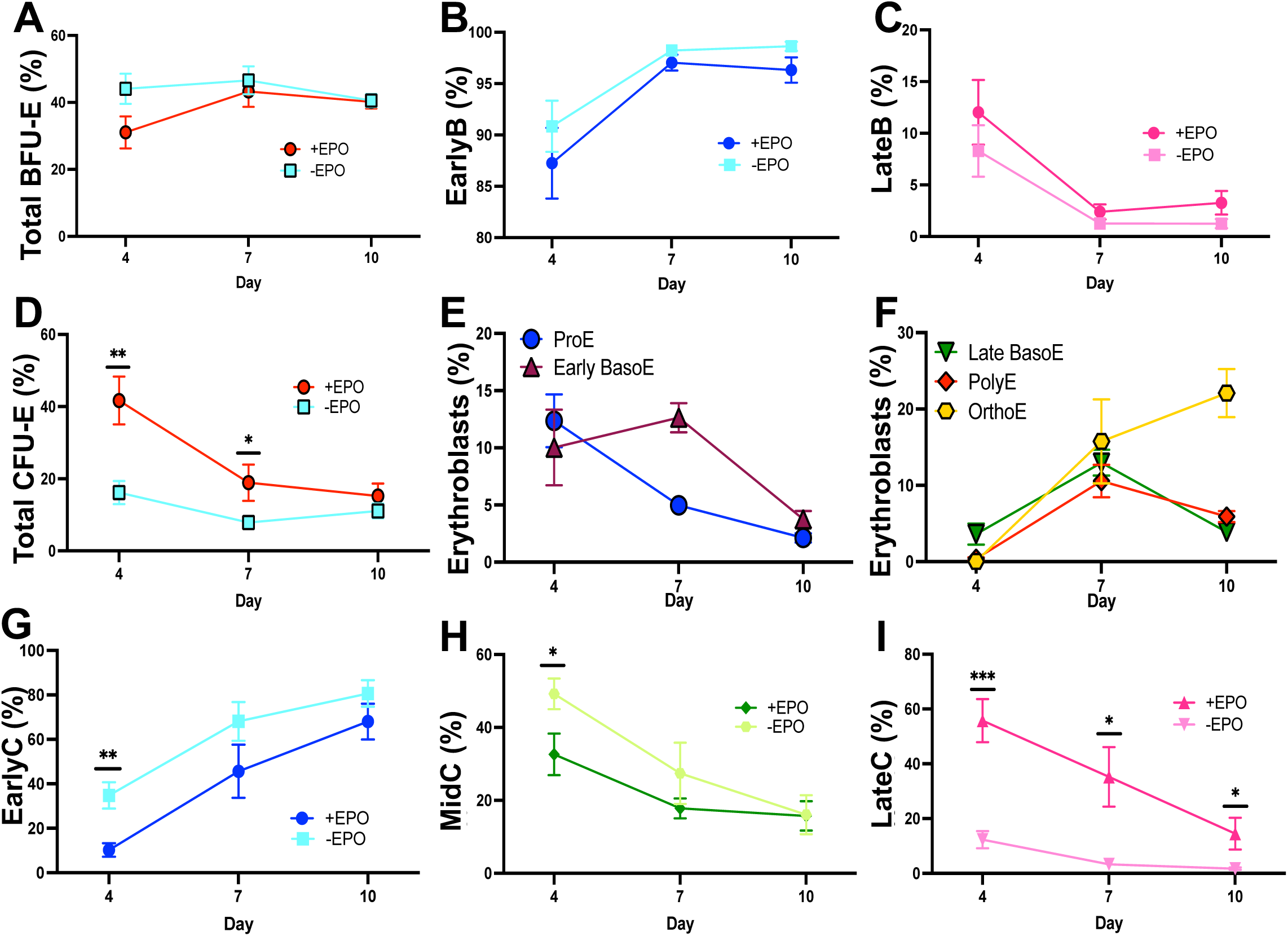
Formation of late CFU-E is dependent on the presence of Epo. Comparison of percentages of the erythroid progenitor and erythroblast populations in BM HSPC cultures incubated for 4, 7, and 10 days in medium with and without Epo. **(A)** Total BFU-E (CD34^+^CD105^+^) are depicted as a percentage of CD49d^+^CD117^+^ cells, **(B)** EarlyB (CD34^+^CD71^lo^CD105^lo^) are depicted as a percentage of total BFU-E, **(C)** LateB (CD34^+^CD71^hi^CD105^lo^) are depicted as a percentage of total BFU-E, **(D)** Total CFU-E (CD34^−^CD105^+^) are depicted as a percentage of CD49d^+^CD117^+^ cells, **(E)** ProE (CD105^hi^CD235a^lo^) and Early BasoE (CD105^hi^CD235a^hi^) are depicted as a percentage of CD235a^+^ cells, **(F)** Late BasoE (CD105^med^CD235a^hi^), PolyE (CD105^lo^CD235a^hi^), and OrthoE (CD105^−^CD235a^hi^) are depicted as a percentage of CD235a^+^ cells, **(G)** EarlyC (CD34^−^CD71^lo^CD105^lo^) are depicted as a percentage of total CFU-E, **(H)** MidC (CD34^−^CD71^hi^CD105^lo^) are depicted as a percentage of total CFU-E, and **(I)** LateC (CD34^−^CD71^hi^CD105^hi^) are depicted as a percentage of total CFU-E. P-values ≤ 0.05 were considered significant (*p < 0.05, **p < 0.01, ***p < 0.001; n=8 for EE populations; n=6 for erythroblast populations).

In previous studies, BFU-Es and CFU-Es have been distinguished based on CD34 and CD36 expression, where BFU-E and CFU-E were characterized as CD34^+^CD36^−^ and CD34^−^CD36^+^, respectively^29,41^. Interestingly, examination by both antibody panels employed in this study showed that CD36 expression within BFU-E and CFU-E populations was unaffected by the absence of Epo (Figures 2D, S2, and S3E-S3F). Further, application of the strategy proposed by Li et al. for analysis of EE populations in +Epo and −Epo cultures failed to resolve Epo-dependent populations (Figure S4A). This is most likely because expression of both CD36 and CD71 is influenced by, but not completely dependent on the presence of Epo, as it is in the case of CD235a (Figures 1D, S1). Yan et al. demonstrated heterogeneity within EE progenitors and proposed strategies for resolution of progenitors and erythroblasts^27,28^. Based on the assessment of CD105 and CD235a, we were able to recapitulate the erythroblast populations within the CD235a^+^ cells (Figure S1C). However, we were unable to separate the BFU-E and CFU-E populations by comparing the patterns of CD34 and CD105 expression (Figure S4B). This could be because a CD34^+^ population that was CD105^hi^ was not observed in our analysis and, thus, the Epo dependence of the subpopulations based on these markers was not discernible.

### All BFU-E and CFU-E Subtypes are Present in Human Bone Marrow

To ascertain that all observed EE subtypes were present in human BM, we performed prospective analyses. For BFU-Es, frozen BM CD34^+^ cells were incubated overnight in pre-stimulation medium, which does not contain Epo, and then analyzed by the immunophenotyping strategy described in Figure 2A. Examination of CD34 expression revealed that >90% of the cells were CD34^+^, suggesting that the short overnight incubation did not result in significant progression beyond the HSPC stages. Immunophenotyping of these cells exhibited the presence of earlyBs and lateBs, with the percentage of lateBs being slightly higher than that seen in BM CD34^+^ cell cultures (Figure S4C). To analyze CFU-E subtypes, total BM mononuclear cells (MNCs) were incubated overnight in pre-stimulation medium and then immunophenotyped as shown in Figure 2A. This MNC analysis revealed the presence of both BFU-E populations and all three CFU-E populations in BM MNCs (Figure S4D). Thus, all EE populations were prospectively observable in BM cells.

### Isolated Early Erythroid Populations Exhibit Expected Differentiation Trajectories

To further investigate the differentiation potential of each of the five EE populations, they were sorted by FACS and recultured in suspension for immunophenotypic analysis and in semi-solid media for CFC assays. BM CD34^+^ HSPCs were cultured and, on day 4, sorted according to the strategy outlined in Figure 2A. In initial attempts, the numbers of earlyCs and midCs were found to be insufficient for downstream analysis, so they were sorted together. For suspension cultures, sorted cells were replated in complete media containing Epo. To examine the cell types that arose from each of the isolated EE populations, immunophenotyping was performed on days 4 and 7 after replating. During reculture, proliferation potential was observed to be highest for earlyBs and lowest for lateCs, while lateBs and earlyCs/midCs exhibited similar intermediate expansion capacity (Figure S4E). On day 4, immunophenotyping of recultured cells from sorted earlyBs detected mostly earlyBs and a few cells belonging to all other EE populations (Figure 4A). By day 7, the sorted earlyBs produced a large number of earlyCs, with a few midCs, lateCs, and erythroblasts as well. Sorted lateBs, on the other hand, produced primarily lateBs, midCs, and lateCs by day 4, which progressed to all erythroblast populations by day 7 (Figure 4B). Notably, earlyCs were not observed in cultures of sorted lateBs, thereby suggesting that lateBs may be transitioning directly to midCs. Reculture of the earlyC/midC pool yielded all CFU-E populations and all erythroblast populations, with mainly ProE and BasoE arising by day 4 and PolyE and OrthoE forming by day 7 (Figure 4C). Sorted lateCs also efficiently produced erythroblasts (Figure 4D). MidCs and lateCs were still present in lateB cultures on day 7, while this was not the case for cultures of earlyC/midCs or lateCs, because all cells had entered TED, as seen in the scatter plots for erythroblasts (Figures 4B, 4C, and 4D). These observations demonstrate that the sorted individual EE populations have high proliferation and differentiation potential and that they are committed to erythroid differentiation. Further, the inability of sorted lateBs to produce earlyCs supports the presence of two differentiation routes.

**Figure 4.**
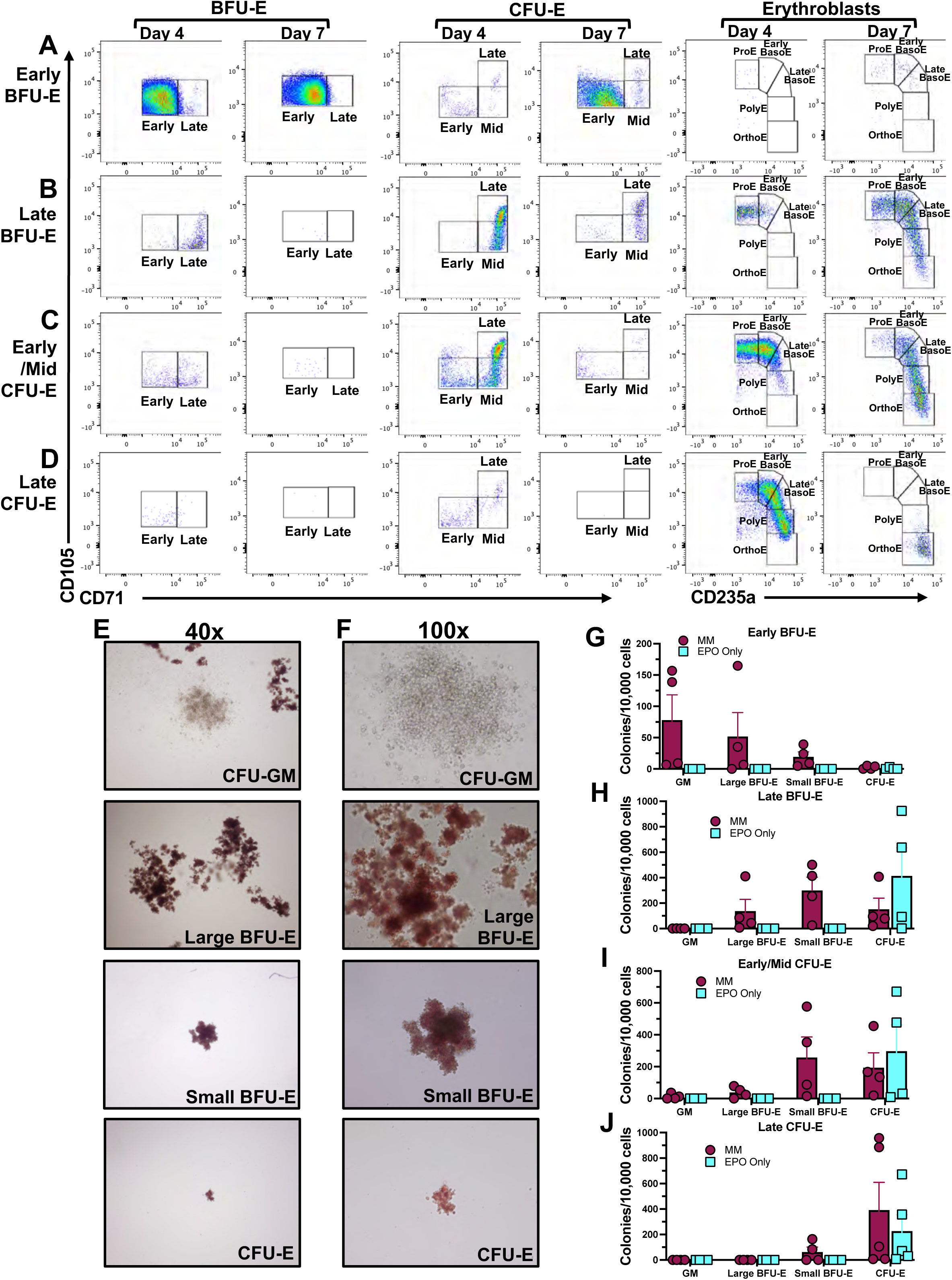
Validation of the erythroid differentiation potential of isolated EE subpopulations. The five EE subtypes were isolated by FACS and then recultured in suspension for immunophenotypic analysis and in semi-solid media for CFC assays. Representative scatter plots for analysis of BFU-E, CFU-E, and erythroblast subpopulations arising from cultures of FACS sorted earlyB (**A**), lateB (**B**), earlyC/midC (**C**), and lateC (**D**) on days 4 and 7 are shown. BFU-E and CFU-E subtypes were analyzed by the immunophenotyping strategy described in Figure 2A, and erythroblasts were analyzed as described by Yan et al. 2021^28^. For CFC assays, sorted cells were cultured in methylcellulose containing SCF, IL-3, GM-CSF, and Epo (MethoCult^®^ classic; MM) or Epo only. Representative images of CFU-GM, large BFU-E, small BFU-E, and CFU-E colonies at 40x (**E**) and 100x (**F**) magnification are shown. Colonies formed by sorted earlyB (**G**), lateB (**H**), earlyC/midC (**I**), and lateC (**J**) were counted on day 14 after plating (n = 4 or 5) and normalized to 10,000 cells. Abbreviations: EPO; erythropoietin, CFU-GEMM; colony forming unit-granulocyte, erythrocyte, monocyte, megakaryocyte, CFU-GM; colony forming unit-granulocyte monocyte, BFU-E; burst-forming unit erythroid, CFU-E; colony-forming unit erythroid, ProE; proerythroblast, BasoE; basophilic erythroblast, PolyE; polychromatic erythroblast, OrthoE; orthochromatic erythroblast, MM; MethoCult^®^ classic with Epo.

For additional functional analysis by CFC assays, the isolated EE cells were plated in semi-solid media containing either all cytokines (including Epo; MM) or Epo alone, as conventionally, CFU-Es have been characterized by their ability to form colonies in Epo-only media^28,29,44^. Analysis of colonies formed by unsorted cells harvested on day 4 of culture revealed the presence of colony-forming unit granulocyte-monocyte (CFU-GM), BFU-E, and CFU-E colonies (Figures 4E, 4F, and S4F). Two types of BFU-E colonies were observed; large colonies that consisted of several clusters of hemoglobinized cells (large BFU-E) and smaller single cluster colonies of a few hundred cells that were visible at 40x (small BFU-E). The CFU-E colonies, on the other hand, were difficult to visualize at 40x, but easily seen at 100x (Figures 4E and 4F). Other studies have also reported such size differences in BFU-E colonies^44,45^.

Examination of colonies formed from the sorted EE populations showed that earlyBs formed CFU-GM or BFU-E colonies in complete medium and did not produce any colonies on Epo-only plates (Figure 4G). LateBs formed both BFU-E colonies and also formed CFU-E colonies in complete medium, and surprisingly, they also produced CFU-E colonies in Epo-alone medium (Figure 4H). While the earlyC/midCs produced small BFU-E and CFU-E colonies in complete medium, they only formed CFU-E colonies in Epo-alone medium (Figure 4I). The last EE population, lateCs, yielded primarily CFU-E colonies in both conditions (Figure 4J). These results suggest that while earlyBs that express CD34 but not CD36 may exhibit some plasticity, retaining the ability to form CFU-GM colonies, the other four EE populations are strictly erythroid lineage committed. They also suggest that lateB and earlyC/midC populations, which form similar types of colonies, represent transitional stages that may be physiologically alike. The lateC represent a subset of CD34^−^CD36^+^ cells that only form CFU-E colonies and have completed the EE transition but have not yet entered TED, as they are CD235a^−^. Overall, analysis by reculturing and CFC assays confirms the potential of the isolated EE progenitor cells to progress along the erythroid trajectory via two parallel routes, one of which transitions through earlyC and the other through lateB (Figure 2F).

### Single Cell RNA-sequencing Identifies Molecular Processes Driving Epo-dependent Transition

To investigate the underlying transcriptional states that drive Epo-dependent transitions within EE populations, we performed scRNA-seq. BM CD34^+^ HSPCs from two healthy donors were cultured for seven days in medium with and without Epo. To avoid perturbations caused by FACS, harvested cells were directly used for the preparation of scRNA-seq libraries using the 10x Genomics platform and sequenced up to a depth of ∼30,000 reads per cell. After sequence mapping, quality control, and normalization, the transcriptomic dataset from all samples were merged in Seurat^46-49^. Unsupervised clustering of the dataset from 26,387 cells yielded 15 clusters (Figure S5A). Assessment of hematopoietic marker genes revealed the presence of clusters specific to HSPCs, megakaryocytes, monocytes, granulocytes, erythrocytes, and early lymphoid progenitors but not mature T and B cells (Figures S5B-S5I). The pattern of canonical human HPSC markers *CD34*, *CD38*, and *PTPRC* (CD45RA) indicated that HSPCs were localized to clusters 0, 9, and 10 (Figure S5B). Expression of lineage markers *IL5RA* (CD125) and *CD14*, mapped granulocytes and monocytes to clusters 4 and 7, respectively, and *GP1BA* (CD42) mapped megakaryocytes to clusters 11 and 12 (Figure S5C-S5E)^49-51^. Furthermore, expression of the early lymphoid genes, *HOPX* and *SPINK2*, mapped to clusters 0, and 9, and the GMP-specific genes, *PRTN3* and *MPO*, mapped to cluster 10 (Figures S5F and S5G)^24,25^. Expression of mature lymphoid markers *CD19* and *CD3D* was observed in a few cells but was not cluster specific and appeared stochastic (Figure S5H). Finally, expression of CD235a identified clusters 6, 3, 1, and 8 as erythroblast-specific (Figure S5I). CD235a^+^ cells represented a large fraction (∼34%) and were aligned with cells derived from cultures containing Epo but absent from cultures lacking Epo (Figure S5J).

To focus on transcriptomic changes associated with erythroid differentiation, we filtered out the non-erythroid lineage clusters (4, 7, 11, and 12), GMP cluster (10) and the two island clusters 13 and 14 that contained only 209 and 158 cells, respectively. Unsupervised reclustering of the remaining subset, consisting of 19,231 cells, generated a second UMAP of eleven clusters (Figure 5A). Pseudotime analysis using Monocle3 provided the progression order of clusters from least to most mature as C0®C1®C9®C2®C3®C4®C10®C6®C8®C5®C7 (Figures 5B and 5C). All clusters were present in +Epo cultures from both donors at comparable percentages, however, clusters C4, C10, C6, C8, C5, and C7 were not observed in −Epo cultures (Figures 5D and 5E). This indicates a clear Epo-dependent transition at the cusp of C3 to C4 transition that coincides with a rise in the expression of CD235a transcripts (Figure 5F).

**Figure 5.**
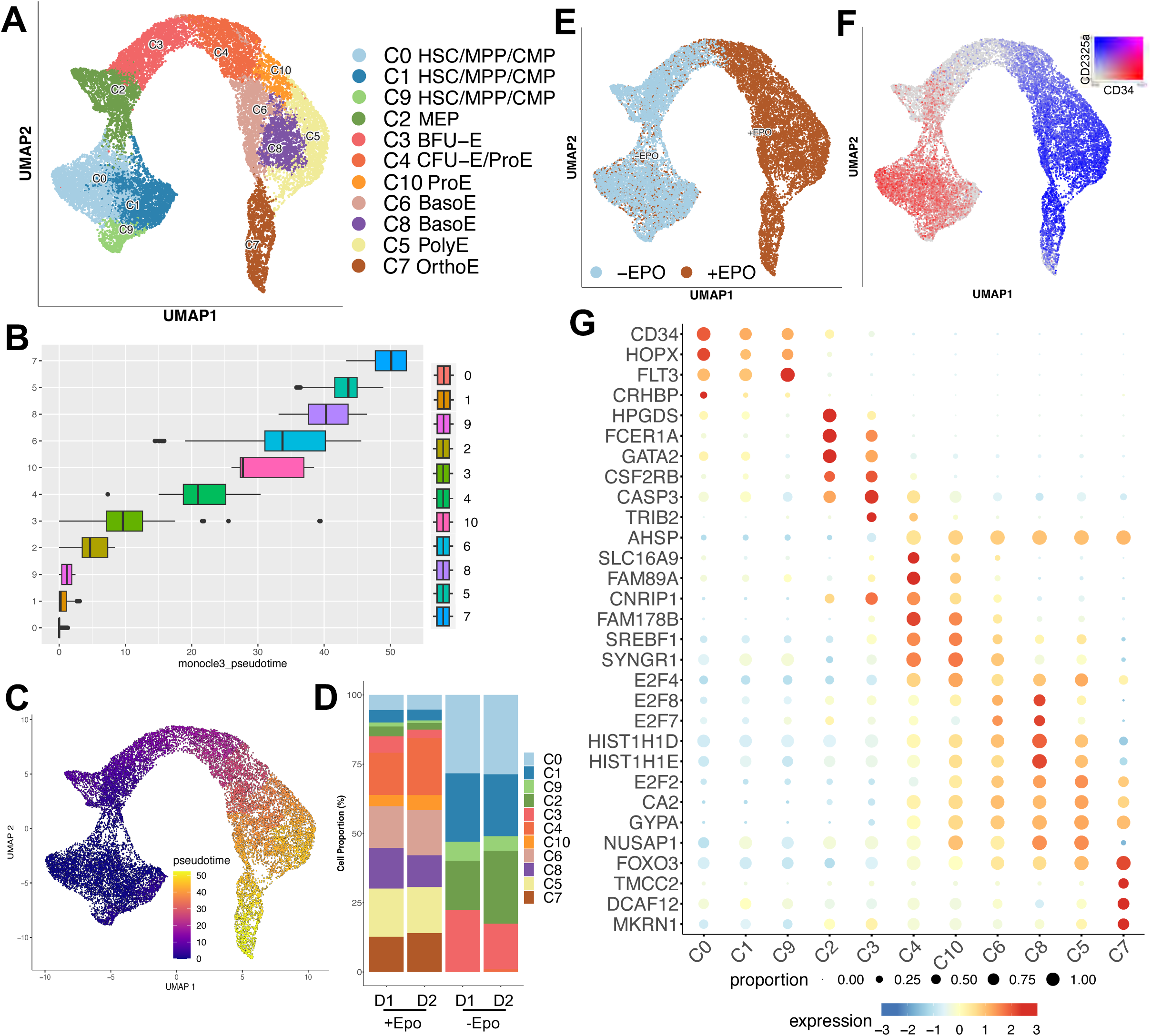
ScRNA-seq analysis reveals an Epo-associated transition. **(A)** UMAP visualization of unsupervised reclustering of HSPC and erythroid cells after filtering out non-erythroid lineages. Cell types are colored by cluster. **(B)** Boxplot depicting the distribution of cells in the psuedotime order of clusters. **(C)** UMAP visualization of cells colored by pseudotime using Monocle 3. **(D)** Bar graphs showing the observed proportion of cells in each cluster for cultures of HSPCs from each donor that were grown in +Epo and −Epo conditions. **(E)** UMAP visualization of cell types involved in erythropoiesis, colored by presence of Epo in culture medium. **(F)** UMAP depicting coexpression of *CD34* and *GYPA* (CD235a). **(G)** Bubble plot depicting expression of representative marker genes used for manual annotation of cell types. Dot size represents the proportion of cells that express each marker, and the filled color represents scaled expression. Abbreviations: HSC; hematopoietic stem cell, MPP; multipotent progenitor, CMP; common myeloid progenitor, BFU-E; burst-forming unit erythroid, CFU-E; colony-forming unit erythroid, ProE; proerythroblast, BasoE; basophilic erythroblast, PolyE; polychromatic erythroblast, OrthoE; orthochromatic erythroblast, Epo; erythropoietin, D1; donor 1, D2; donor 2.

We used the pseudotime order of clusters, along with the expression of previously described representative stem, progenitor, and erythroblast genes, for manual annotation of the eleven clusters (Figures 5G and S6, and Table S2)^24,25,29,52-55^. C0, C1, and C9 exhibit expression of several HSC, MPP, and CMP markers, which, in addition to *CD34*, include *HOPX*, *FLT3*, and *CRHBP*. Cluster C2, on the other hand, expresses several MEP genes, including *HPGDS*, *FCER1A*, and *GATA2*, but does not express *FLT3*. HSPC surface markers that are commonly employed for immunophenotyping, including *CD34*, *THY1* (CD90), *Flt3* (CD135), *CD38, IL3RA* (CD123), *PTPRC* (CD45RA), and *KIT* (CD117), were all expressed in C0, C1, and C9 that were annotated as HSC/MPP/CMP (Figure 5A and S6B). Transcripts for *PTPRC* (CD45RA), *CD38*, and *KIT* (CD117) were also expressed in MEP cluster C2. There were two surprising observations in our dataset. First is the downregulation of *CD34* transcription from C2 onwards; surface expression of CD34 protein on MEPs and BFU-Es is well documented and observed in our experiment as well. Second is the presence of *IL3RA* (CD123) transcript in MEP containing C2, whereas the protein is normally not observed^56^.

To annotate BFU-Es, CFU-Es, and erythroblast clusters, we examined the expression of representative marker genes that have been reported in these populations or described as early erythroid genes in previous RNA-seq analyses^24,25,29,52^. BFU-E genes *CSF2RB*, *CASP3*, and *TRIB2* are highly expressed in C3, whereas CFU-E-specific *AHSP* arises in C4 (Figure 5G and Table S2). Transcripts for three other CFU-E genes, *SLC16A9*, *FAM89A*, and *CNRIP1*, are also high in C4. These patterns suggest that C3 and C4 contain BFU-Es and CFU-Es, respectively. Significant upregulation of *GYPA* (CD235a) and *SLC4A1* (CD233) suggests the presence of erythroblasts in C4, C10, C6, C8, C5, and C7 (Figure S6C). ProE-specific genes *FAM178B*, *SREBF1*, *SYNGR1*, and *E2F4* are high in C4 and C10, thereby suggesting that ProEs may be distributed between these two clusters. Similarly, examination of BasoE genes, *E2F8*, *E2F7*, *HIST1H1D*, and *HIST1H1E* suggested their distribution between C6 and C8. Genes for PolyE (*E2F2*, *CA2*, *GYPA*, and *NUSAP1*) and OrthoE (*FOXO3*, *TMCC2*, *DCAF12*, and *MKRN1*) were highly expressed in C5 and C7, respectively. Thus, based on these expression patterns, the eleven clusters were annotated in order of pseudotime: C0, C1, and C9 as HSC/MPP/CMP, C2 as MEP, C3 as BFU-E, C4 as CFU-E and ProE, C10 as ProE, C6 and C8 as BasoE, C5 as PolyE, and C7 as OrthoE (Figure 5A and 5G).

Analysis of differentially expressed genes (DEG) in all eleven clusters using Seurat revealed additional cluster specific genes and Kyoto Encyclopedia of Genes and Genomes (KEGG) pathways underlying the successive transitions along erythroid differentiation (Figure 6). Expression of the top 20 differentially upregulated genes within C0, C1, and C9 indicate overlapping profiles (Figure 6A and Table S3). Among clusters spanning erythroblast stages, the transcriptional profile of C10 exhibits similarity with C4 and C6, and that of C8 exhibits similarity with C6 and C5. These patterns indicate a gradual activation and repression of stage-specific genes, suggesting an erythroblast continuum. OrthoEs, however, form a distinct cluster (C7) and exhibit a unique expression profile.

**Figure 6.**
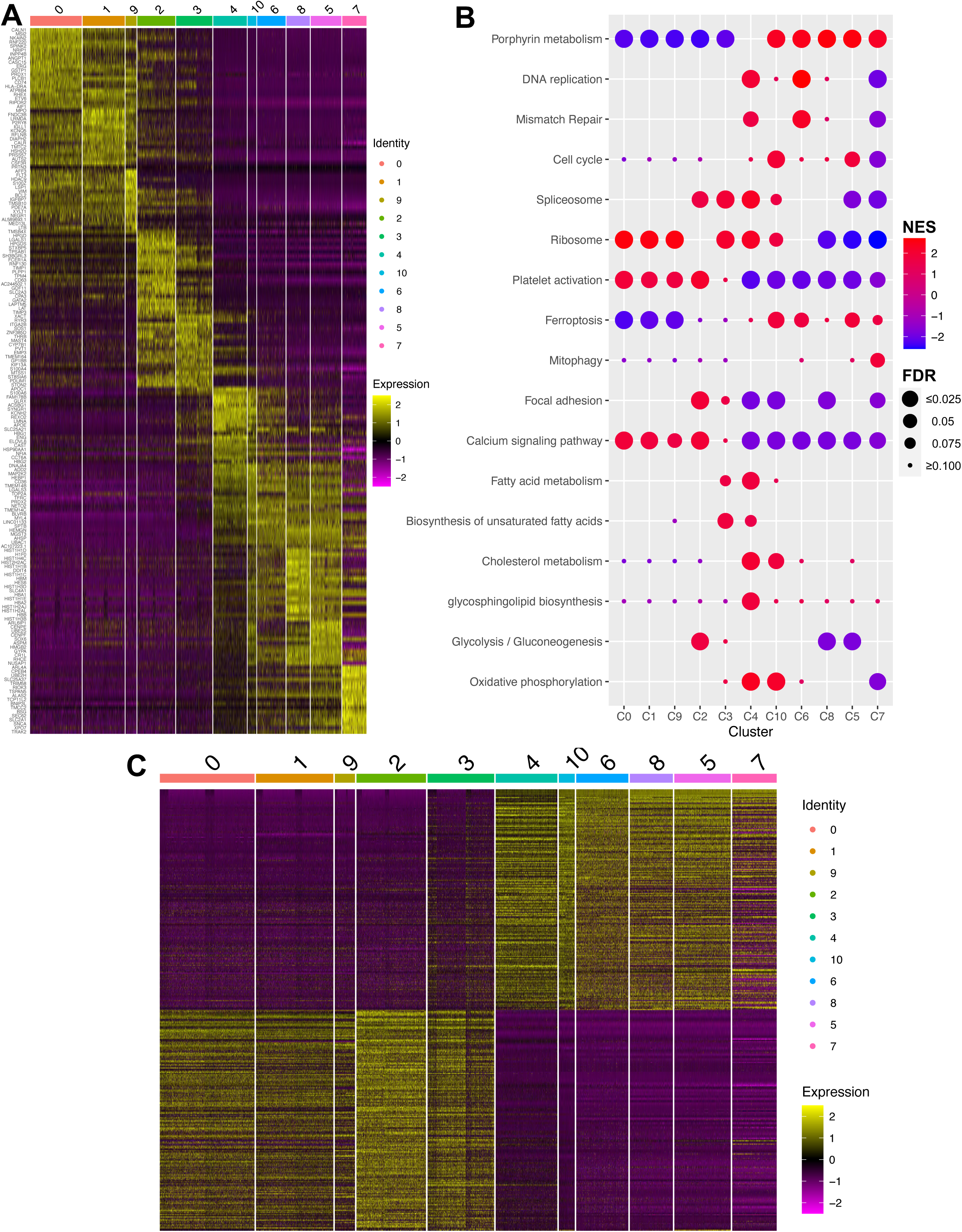
Differentially expressed genes and gene ontology analysis reveal Epo-dependent genes and cellular pathways. **(A)** Heatmap depicting expression trends of the top 20 differentially upregulated genes in each cluster. Columns are ordered according to pseudotime showing progression from immature to mature cells. **(B)** Bubble plot of terms found to be enriched in gene set enrichment analysis of each cluster. The dot color represents the normalized enrichment score (NES), and the size indicates the false discovery rate (FDR). **(C)** Heatmap depicting expression trends for the top 200 (up- and downregulated; 400 total) Epo-dependent DEGs across all clusters. Columns represent clusters ordered according to pseudotime.

To identify functional differences between clusters, we performed gene set enrichment analysis (GSEA)^57^. For this, the entire gene set for each cluster was used as an input into GSEA, and predominant KEGG pathways were assessed across all clusters (Figure 6B). This analysis confirmed the involvement of several well-established erythroid differentiation pathways. Top amongst these is protoporphyrin metabolism, which is upregulated from C10 onwards. Upregulation of DNA replication, mismatch repair, and cell cycle genes is in alignment with the known expansion of erythroblasts during TED. Similarly, upregulation of RNA processing complexes—spliceosome and ribosome—may be needed to meet the needs of proliferating progenitors and erythroblasts. All of these processes, except protoporphyrin metabolism, decline later in the erythroblasts preceding enucleation. Other expected observations include a downregulation of platelet activation and a slight upregulation of ferroptosis and mitophagy in erythroblast clusters. Events that have been previously reported to have a role in erythropoiesis but not well characterized include focal adhesion and calcium signaling, both of which were downregulated^58,59^. Very interestingly, we found transient upregulation of lipid and cholesterol metabolism pathways. Biosynthesis of unsaturated fatty acids was enriched in C3 and C4, cholesterol metabolism was up in C4 and C10, and glycosphingolipid biosynthesis was up in C4 (Figure 6B). Other interesting metabolic events include a transient reduction in glucose metabolism and oxidative phosphorylation in late erythroblast-specific clusters (C8, C5, and C7), which have recently been reported to occur in mouse and human erythroblasts in other studies as well^60-62^. Overall, the scRNA-seq and GSEA analyses revealed transcriptional alterations across all erythroid differentiation stages and identified potential involvement of lipid metabolic pathways in Epo-dependent transitions.

### Differentially Expressed Genes Reveal Epo-associated Transcriptional Dynamics

Next, we wanted to assess transcriptional changes associated with the Epo-induced C3 to C4 transition. For this, C3 and C4 were subset, and DEG analysis was then performed based on Epo treatment (Table S4). This identified 1939 Epo-dependent genes, of which 695 were upregulated and 1244 were downregulated (. Assessment of the top 400 Epo-dependent genes (200 upregulated and 200 downregulated) revealed a switch in expression pattern between clusters prior to the onset of Epo-dependence (C0, C1, C9, C2, and C3) and those that follow (C4, C10, C6, C8, C5, and C7) (Figure 6C). The Epo-dependent DEG list includes previously reported genes such as the three erythroid master transcription factors (*GATA1*, *TAL1*, and *KLF1*), genes involved in heme and hemoglobin synthesis (*HBA2*, *HBA1*, *AHSP*, *HBG1*, *HBB*, etc.), and several others such as *EPOR*, *ERFE*, *HES6*, and *JAK2*. Upregulation of these genes was observed in cells grown in presence of Epo and in clusters that arise after the C3 to C4 transition (Figure S7A and S7B). Most importantly, our analysis identifies several genes that were so far not known to be Epo-induced (Table S4). Interesting amongst these are genes involved in lipid and cholesterol metabolism and significantly upregulated at or after the onset of Epo-dependent transition. Some of these genes, including *NCEH1*, *SLC25A21*, and *TLCD4*, arise in CFU-Es (C4) and continue to be expressed in erythroblasts (Figure S7C). Others, including *ACSBG1*, *LRP1*, *APOC1*, and *APOE*, are transiently expressed, arising in either C3 (BFU-E) or C4 (CFU-E) and downregulated in erythroblasts (Figure S7D). Still others, such as *OSBP2*, *TSPO2,* and *PHOSPHO1*, that have previously been shown to be critical for TED in mice, were also detected and were highly expressed in clusters C8, C5, and C7 that are specific to late BasoE, PolyE, and OrthoE, respectively (Figure S7E)^63,64^. Thus, the scRNA-seq analysis allows assessment of Epo-associated and stage specific transcriptomic patterns and identifies several new Epo-induced genes.

### Epo-induced High Coexpression of CD71 and CD105 in the CFU-E Cluster

While cluster annotation placed BFU-Es and CFU-Es in clusters C3 and C4, respectively, we wanted to determine the potential location of the five EE populations and check if the single cell transcriptomic analysis recapitulates the pattern of CD36 and the Epo-associated high coexpression of CD71 and CD105 in the CFU-E containing cluster C4. For this, we compared CD71, CD105 and CD36 expression within clusters and examined the percentage of cells coexpressing these markers (Figure 7A-7D). CD71 and CD105 have previously been observed on MEPs; however, in our analysis, in addition to C2, low levels of their transcripts were also present in C0, C1, and C9 (Figures 7A and 7B). CD71 was upregulated from C3 onwards, with expression levels maintained through C5 (Figure 7A). Since almost 100% of C3 cells express CD71, lateBs and midCs are likely present in C3 (Figure 7C). CD105 upregulation, on the other hand, was transient, being highest in C4 and downregulated sharply afterwards (Figure 7A and 7C). The highest expression of CD71 and CD105 in C4 combined with largest percentage of cells co-expressing these two markers in C4 suggests that CD71^hi^CD105^hi^ lateCs are located within C4 (Figures 7A and 7D). Although CD36 lagged CD71, arising in C2, its expression largely paralleled that of CD71; they were coexpressed in almost 100% of the cells from C4 to C5, and downregulated in C7 (Figure 7D). The highest coexpression of CD36 and CD105 also in C4 (∼96% of cells) is in alignment with the location of lateCs in this cluster, as lateCs were ∼100% CD36^+^ by immunophenotyping (Figures 7D and 2E). The presence of an intermediate percentage of cells expressing CD36 in C3 (∼75%) also suggests that lateB and midC are likely in this cluster (Figure 7C). Overall, the Epo-dependence of the C4 cluster and the highest coexpression of CD71, CD36, and CD105, even at the level of transcription, in C4 support our finding that Epo is required for the acquisition of the CD71^hi^CD105^hi^ immunophenotype exhibited by lateCs.

**Figure 7.**
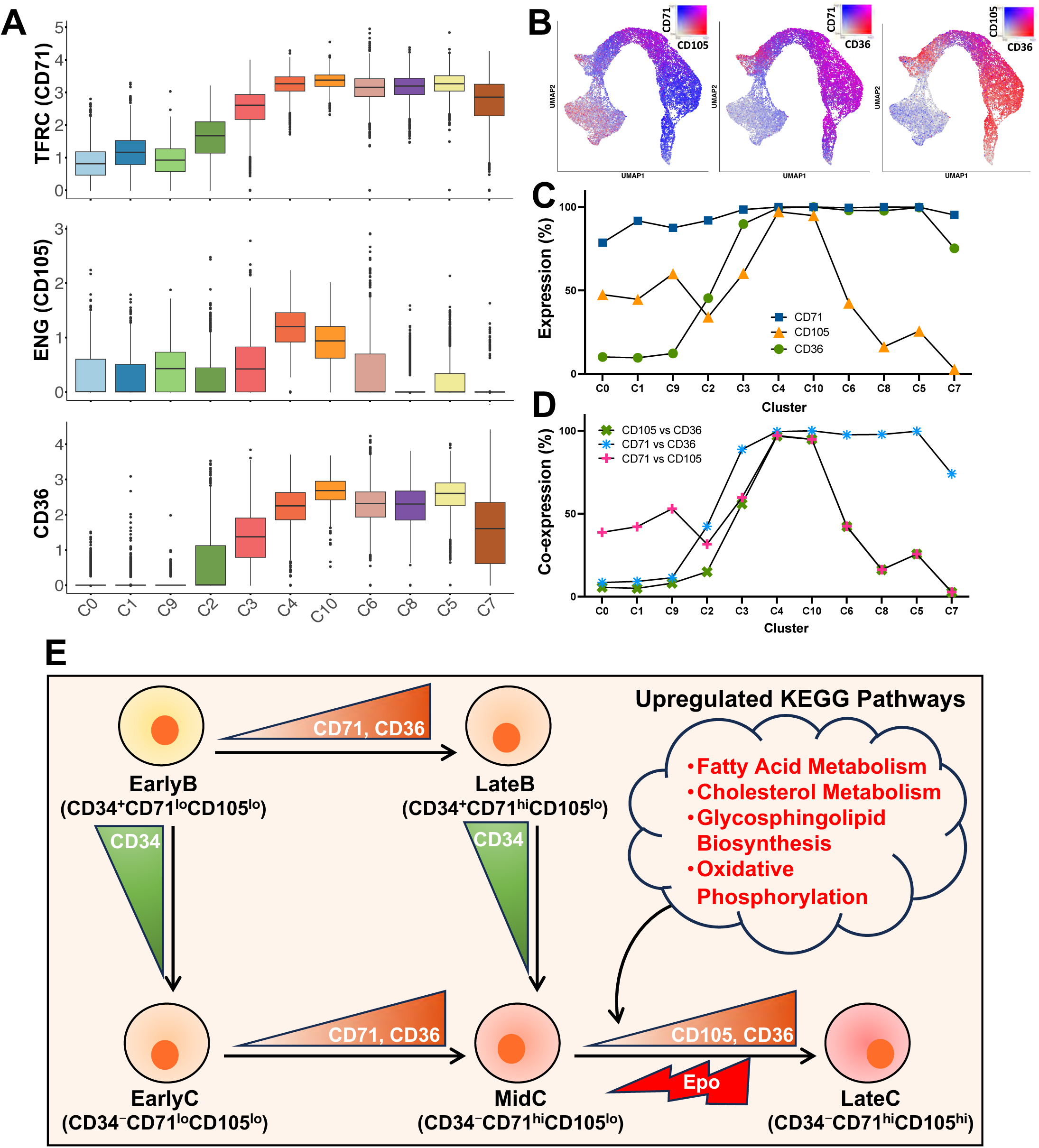
Coexpression of CD71, CD105, and CD36 in lateC cluster C4. **(A)** Boxplots demonstrating the expression of CD71, CD105, and CD36 transcripts in each cluster. **(B)** UMAP plots for coexpression of CD71 and CD105, CD71 and CD36, and CD36 and CD105 transcripts. **(C)** Graph depicting percentage of cells expressing CD71, CD105, and CD36 transcripts in each cluster. **(D)** Graph depicting percentage of cells coexpressing CD71 and CD105, CD105 and CD36, and CD71 and CD36 transcripts in each cluster. **(E)** Schematic representation of the two routes to formation of lateC. One passes through the earlyC stage, whereas the other bypasses earlyCs, directly forming midCs that transition to the Epo-dependent CD34^−^CD71^hi^CD105^hi^ lateC cells. EarlyB to lateB and earlyC to midC transitions are accompanied by increase in expression of CD71 and gain of CD36 positivity, whereas earlyB to lateB and earlyC to midC transitions are accompanied by loss of CD34 positivity. KEGG pathways upregulated during Epo-dependent converstion of midC to lateC are shown. Abbreviations: Epo; erythropoietin, earlyB; early BFU-E, lateB; late BFU-E, earlyC; early CFU-E, midC; mid CFU-E, lateC; late CFU-E.

## Discussion

Commitment to erythroid differentiation occurs during EE, and thus, elucidation of the critical role of Epo in driving the development of BFU-E and CFU-E and their transition to ProE is crucial for advancing the knowledge of underlying regulatory mechanisms. By immunophenotyping transiently expressed erythroid markers on cells from *ex vivo* cultures of BM-derived HSPCs, we have been able to resolve heterogeneous EE progenitor populations and identify the key Epo-induced transition (Figure 7E). Based on the expression patterns of CD34, CD71, and CD105 within Lin^−^CD123^−^CD235a^−^CD49d^+^CD117^+^ cells, we detect five EE subtypes: two BFU-E and three CFU-E. The two BFU-Es, earlyB and lateB are CD34^+^ but can be distinguished as CD71^lo^CD105^lo^ and CD71^hi^CD105^lo^, respectively. The CFU-Es earlyC, midC, and lateC, are CD34^−^ and distinguishable as CD71^lo^CD105^lo^, CD71^hi^CD105^lo^, and CD71^hi^CD105^hi^, respectively. Additionally, prospective analysis detected the two BFU-E subtypes and three CFU-E subtypes in BM cells, and after isolation, each individual population exhibited the capacity to continue differentiation along the erythroid route. Importantly, Epo was found to be essential for the acquisition of the CD71^hi^CD105^hi^ immunophenotype that accompanies the midC to lateC transition. The lateC is a previously undescribed unique Epo-dependent population that is CD235a^−^ but expresses high levels of CD71, CD105, and CD36. Expression pattern of these markers allowed us to decipher two routes to lateC formation (Figure 7E). In the first, loss of CD34 expression occurs prior to gain of CD36 and progresses from earlyB➔earlyC➔midC➔lateC. The second pathway, on the other hand, bypasses earlyCs because CD36 expression begins in cells still expressing CD34, and progression follows from earlyB➔lateB➔midC➔lateC.

Although not entirely Epo-dependent, both CD71 and CD105 are dynamically expressed and sensitive to Epo in EE cells. While the function of CD71 (transferrin receptor) is well characterized, the role played by CD105 (endoglin) in erythroid differentiation is less clear^65,66^. Also known as transforming growth factor β (TGF-β) receptor type III, CD105 associates with type I and II receptors upon binding of TGF-β superfamily ligands to regulate gene expression via Smad signaling^67-69^. In mice, CD105 loss was reported to be associated with decreased erythroid output and a potential role in regulating proliferation of BFU-Es and BasoEs^70,71^. Expression of CD105 on immature human erythroid cells has been known for a long time, and its increase in MEPs has been positively corelated with erythroid commitment^32,72^,. We observed a basal level of CD105 transcripts within HSPC clusters (this study) and surface proteins on MEPs, CMPs, and GMPs by flow cytometry (Schippel and Sharma, unpublished data). Importantly, we see an Epo-dependent transient upregulation of CD105 mRNA and protein during EE. Further characterization of CD105’s function is needed to identify its target genes and cellular processes in EE cells.

In several studies, expression of CD36 has been proposed to mark commitment to the erythroid lineage^30-33,35,73^. The onset of CD36 expression has been shown to be accompanied by the loss of CD34 during the BFU-E to CFU-E transition^29,36^. Studies on the differentiation of human peripheral blood (PB) CD34^+^ cells have identified a CD34^+^CD36^+^ population that gives rise to BFU-Es and CFU-Es^27^. Accumulation of a CD34^+^CD36^+^ population has also been detected in the PB of patients with hypercortisolemia and is associated with unresponsiveness to dexamethasone, a glucocorticoid commonly used for anemia treatment^36,74,75^. In our study, the CD34^+^CD36^+^ lateBs were identified as an intermediate population that forms in an Epo-independent manner and transitions directly to the CD34^−^CD36^+^ midC. Interestingly, humans lacking CD36 protein due to genetic mutations generally develop normally, but have thrombocytopenia, increased bleeding, and RBC abnormalities such as higher RBC distribution width (RDW)^76-78^. CD36 knockout mice also develop normally, although they also have thrombocytopenia, along with slight but significantly reduced RBC counts, hemoglobin, and hematocrit levels, and a 40% decrease in the colony-forming ability of their HSCs^76,79^. In contrast, in *ex vivo* experiments, CD34^+^ HSPCs from an individual lacking CD36 were found to undergo erythroid commitment and differentiation without much effect on either CD71 and CD235a expression or erythroblast development, as assessed by CD49d (α-4 Integrin) and CD233 (Band 3) expression using the strategy proposed by Hu et al.^30,80^. However, whether CD36 loss affects other markers such as CD105 or the enucleation and maturation of the erythroblasts remains to be determined.

In most previous studies analyzing EE cells and erythroblasts, BM-, PB-, or cord blood (CB)-derived CD34^+^ HSPCs were cultured in the presence of Epo, along with other cytokines^27-30^. Our approach of comparing +/−Epo conditions facilitated segregation of cell types in an Epo-dependent manner. Profiling of the Epo-associated transcriptomic changes across all stages of erythroid differentiation complemented our immunophenotypic analysis and identified C4 as the cluster that exhibits the highest coexpression of CD71, CD105, and CD36 and harbors lateCs. In alignment with many published transcriptomic and proteomic analyses, our single cell analysis detected previously reported Epo-dependent pathways and genes, and, importantly, enabled the detection of new Epo targets. Notable amongst these are the genes involved in lipid and cholesterol metabolism, the significance of which is underscored by the numerous studies that demonstrate a link between these metabolic processes and erythroid differentiation in mice, humans, and other species^60,63,64,81-84^. Defects in lipid uptake are associated with impairment of RBC maturation in mice^85^. Additionally, chronic anemia patients with high erythropoietic activity have been reported to have hypocholesterolemia^86^. Although *de novo* synthesis has been shown to be downregulated during TED, maintenance of cholesterol levels was found to be critical for the integrity and lifespan of RBCs, thereby indicating the importance of cholesterol uptake and accumulation^60,87,88^. We observed a transient expression of lipid homeostasis genes (*ACSBG1*, *LRP1*, *APOC1*, and *APOE*) in BFU-E and CFU-E clusters and a continued expression of others into TED (*NCEH1*, *SLC25A21*, and *TLCD4*). These observations are supported by a recent study that shows an association between lipoprotein metabolism and erythroid potential of HSCs, specifically demonstrating ApoE-mediated enhanced erythropoiesis^89^. Functional characterization of these newly identified Epo-dependent genes will reveal the hitherto undefined role of lipid metabolism in the regulation of EE in humans.

Anemia is a common health condition that can arise due to a multitude of health conditions, including iron or vitamin deficiency, inherited RBC disorders, infections (such as malaria), obstetric-gynecological conditions, and chronic diseases that lead to dysregulated erythropoiesis^90,91^. Administration of recombinant human Epo and dexamethasone are prevalent treatments for anemia; however, they are not always effective, and the observed improvements are often temporary^92,93^. Thus, there is a significant need to advance the knowledge of Epo’s influence on downstream cellular processes. Our study provides a framework for examining cellular pathways and molecular processes controlled by Epo. This approach can be applied for investigations into regulation by other factors that may promote erythroid differentiation to aid the identification of new therapeutic targets.

## Materials and Methods

### Culture and Differentiation of CD34^+^ HSPCs

Human BM-derived CD34^+^ HSPCs and MNCs were purchased from Ossium Health (San Francisco, CA). Quality control reporting >90% viability and >90% CD34 positivity was provided for all donor samples. MS-5 cells (used as feeder cells for co-culture with CD34^+^ HSPCs) were maintained in Dulbecco’s modified Eagle’s medium (DMEM) with high glucose and L-glutamine (Thermo Fisher Scientific, Waltham, MA), 10% fetal bovine serum (FBS, Omega Scientific, Tarzana, CA), and 1% penicillin-streptomycin (Pen-Strep, Sigma-Aldrich, St. Louis, MO).

Culture of BM CD34^+^ HSPCs was performed as previously described^37-39^. Briefly, on day 0, HSPCs were plated at a density of 1 x 10^5^ cells/well of a 48-well plate in 200 μl of pre-stimulation medium (serum-free X-VIVO 15 containing 2 mM L-glutamine, 50 ng/mL stem cell factor (SCF), 50 ng/mL thrombopoietin (TPO), 20 ng/mL interleukin-3 (IL-3), and 50 ng/mL Flt3-Ligand (Flt-3)). On day 1, the pre-stimulated CD34^+^ cells were collected and plated onto MS-5 cells at a density of 3 x 10^4^ cells/well for analysis on day 4 or 1 x 10^4^ cells/well for analysis on day 7 or 10) in 200 μl of myeloid medium (MM) (DMEM with high glucose and L-glutamine, 10% FBS, and 1% Pen-Strep, supplemented with 50ng/ml Tpo (Miltenyi Biotec, Auburn, CA), 5ng/ml SCF, 5ng/ml IL-3, and 5ng/ml Flt-3, with or without 4U/ml Epo (all BioBasic, Amherst, NY). Half-media changes were performed on days 4 and 7.

For reculture experiments, the individual EE cells were sorted into complete media (containing Epo) supplemented with 20% FBS as described below. The sorted cells were counted and plated on MS-5 cells at a density of either 3 x 10^4^ cells/200 μl or 1 x 10^4^ cells/200 μl and incubated as described above.

### Immunophenotyping and Cell Sorting

For flow cytometry analyses, cells were harvested and stained with fluorescently labeled antibodies as previously described^37-39^. Briefly, cells were washed in phosphate-buffered saline (PBS), counted using the Countess 3-FL Automated Cell Counter (Thermo Fisher Scientific, Waltham, MA), resuspended in PBS, and then stained with an antibody cocktail, as indicated in the figures. Stained cells were washed with PBS, fixed with 1% paraformaldehyde (PFA) (Sigma Aldrich, St. Louis, Missouri), and resuspended in fluorescence activated cell sorting (FACS) buffer (PBS with 2% FBS and 5 mM EDTA). Single stained controls were prepared using compensation beads (UltraComp eBeads from Thermo Fisher Scientific, Waltham, MA) as per the manufacturer’s recommendations. All flow cytometry analysis was performed on a Canto II (BD Biosciences, San Jose, CA), and data was analyzed using FlowJo (version 10.9.0; flowjo.com). Bar and line graphs were generated, and statistical analysis by student T-test was performed in GraphPad Prism (version 10.0.3).

For FACS, cells were collected on day 4 of culture, washed, counted, and stained as described above, but not fixed, and then resuspended in FACS buffer at a density of 3-5 x 10^6^ cells/ml. Cells were sorted into four populations (earlyB, lateB, earlyC/midC, and lateC) based on the strategy outlined in Figure 2A using a FACS Aria II (BD Biosciences, San Jose, CA). Sorted cells were counted and plated either in complete myeloid medium for re-culture or in semi-solid medium for CFC assays.

### Colony Forming Cell Assay

For CFC assays, MethoCult^®^ classic (H4434), classic without Epo (H4534), Epo-only (H4330), or without cytokines (H4230) were used (STEMCELL Technologies; Cambridge, MA). Cells were diluted to a density of 2.5–5 x 10^3^ cells/ml in medium and plated in triplicate in 35 mm tissue culture dishes. After 14 days of incubation, colonies were counted and phenotyped based on criteria described by Palis and Koniski^44^. Images were obtained using an Olympus IX2-SP inverted microscope under 40x and 100x magnification.

### Single Cell RNA Sequencing

Cells were collected from human BM CD34^+^ HSPCs cultures grown in myeloid medium with and without Epo for seven days, washed and resuspended at a density of 1 x 10^3^ cells/ml in ice-cold 1x PBS. Libraries were prepared using the Chromium Next GEM Single Cell 3’ Reagent Kit v3.1 (Dual Index; 10x Genomics) according to the manufacturer’s instructions. Briefly, cells were partitioned into gel beads by running them through the Chromium iX controller (10x Genomics). Full-length cDNA was generated using the 10x Genomics platform in a Bio-Rad C1000 thermal cycler. Gel beads-in-emulsion (GEMs) were cleaned with Dynabeads cleanup mix (see manufacturer’s protocol), and cDNA was then amplified by PCR. Amplified cDNA was cleaned with solid-phase reversible immobilization (SPRI) beads and then quantified using a Qubit^®^ 2.0 fluorometer (ThermoFisher; Waltham, MA). For the preparation of 3’ gene expression dual index libraries, cDNA fragmentation, end repair, and A-tailing were performed, and then size selection was done using SPRI beads (10x Genomics). Oligonucleotide adaptors were ligated, dual index TT (set A) primers were incorporated using PCR, and DNA was again quantified on a Qubit^®^ 2.0 fluorometer. Finally, the libraries were sequenced using the Novogene platform (paired-end sequencing, 150 bp reads; PE150).

### Single Cell Transcriptomic Data Analysis

After sequencing, Cell Ranger v7.1.0 (10x Genomics) was used for the processing of raw sequencing data (FASTQ files), mapping reads to the genome, and generation of unique molecular identifier (UMI) counts for gene expression via gene-barcode matrix^94^. The software package Seurat version 4.4.0 was used to import data for all four samples, only considering features observed in more than three cells^95^. Additional quality control was performed, including filtering out cells with less than 500 features (low quality or cell fragments), greater than 7500 features (doublets), or >10% of reads mapping to the mitochondrial genome. Expression levels were then log normalized to a factor of 10,000. The top 2000 variable features were identified using the *FindVariableFeatures* function. The data was then scaled (*ScaleData* function), and principal component analysis (PCA) was performed (*runPCA* function). Unsupervised clustering was performed by applying the *FindNeighbors* and *FindClusters* functions. UMAPs were then generated using *runUMAP*. Finally, doublets were identified, using the DoubletFinder package in R, and subsequentially filtered out in Seurat^96^. This process was repeated for all four samples (Donor 1 and Donor 2 in +/−Epo conditions; resolution 0.5), and then the samples were merged using the *Merge* function in Seurat. Unsupervised clustering was performed again on the merged object (resolution 0.4), and ShinyCell and Seurat were used to generate UMAPs, boxplots, graphs, and heatmaps^97^. Clusters containing cell types not involved in erythropoiesis, as well as island clusters containing less than 500 cells, were filtered out using the *subset* function. The remaining cells were re-clustered as described above (resolution 0.5). Pseudotime analysis was performed using Monocle3^98,99^ and DEG analysis were performed using *FindAllMarkers* (Seurat). For DEG analysis comparing clusters C3 and C4 by Epo treatment, a false discovery rate (FDR) <0.05 and log fold change (Log2FC) of ±0.2 (20% or larger change in expression) was considered significant. Gene set enrichment analysis (GSEA) analysis was performed using the WEB-based GEne SeT AnaLysis Toolkit (WebGestalt) platform; a false discovery rate (FDR) <0.05 was considered significant^100^. All relevant software and packages, along with their versions, can be found in Table S5.

## Acknowledgments

This work was supported by funds to SS from the National Institutes of General Medical Sciences (GM127464) and National Cancer Institute (P30CA023074) of the National Institutes of Health, the Valley Research Partnership Program (VRP P1-4009 and VRP77), and the Arizona Biomedical Research Center (ABRC: RFGA2022-010-30). The content is solely the responsibility of the authors and does not necessarily represent the official views of the National Institutes of Health.

---

**Figure S1.**
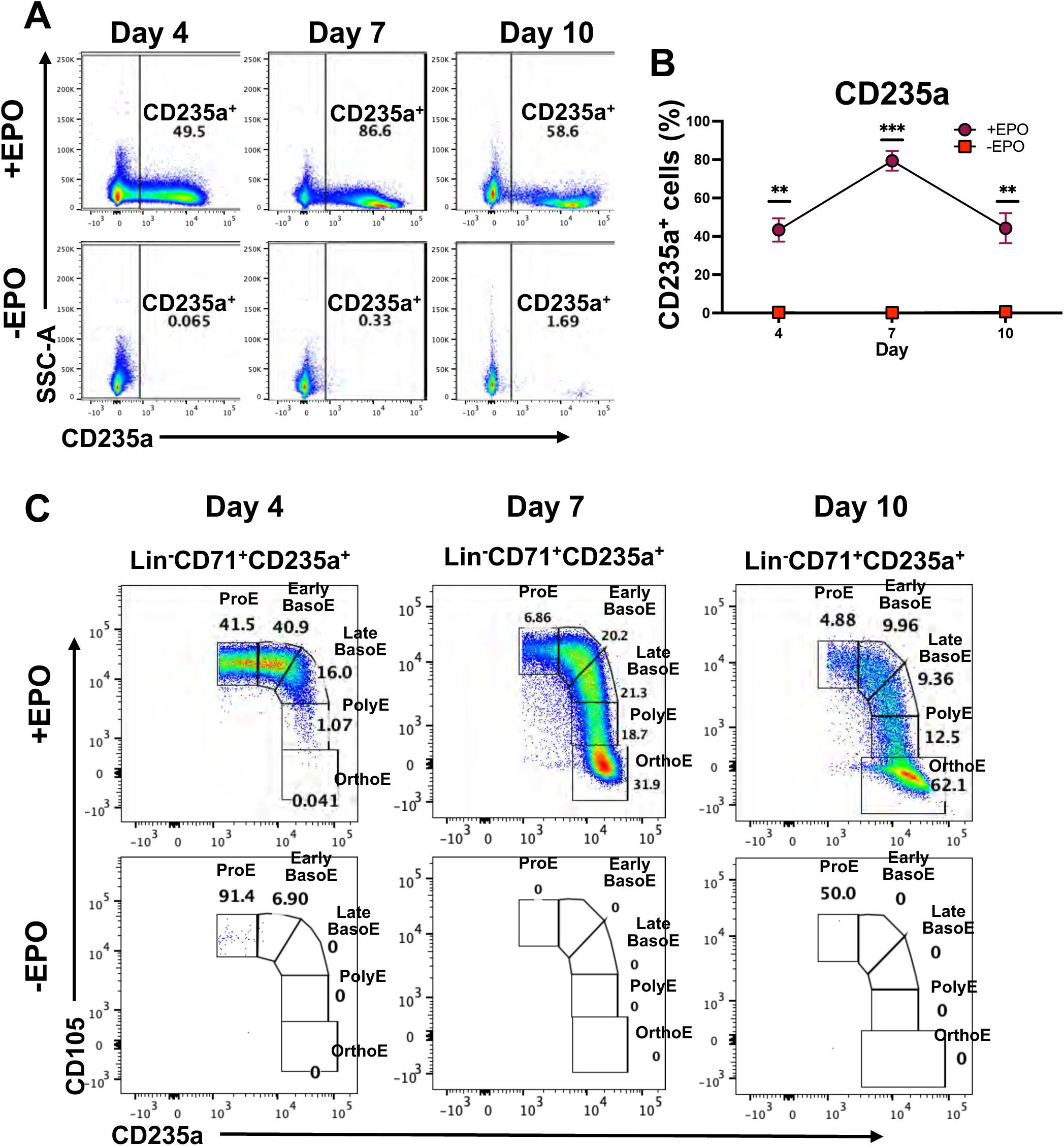
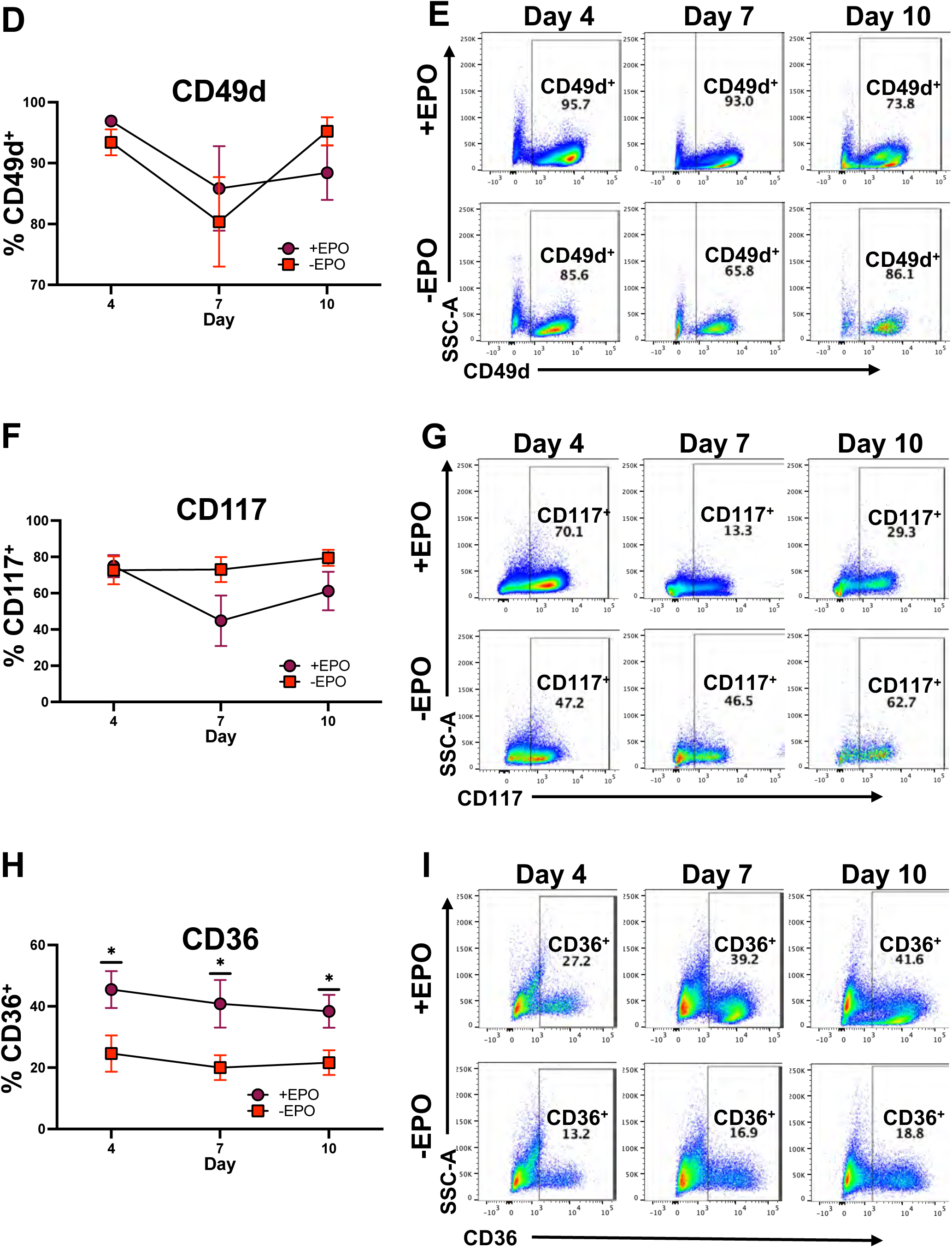
Epo-dependence of early erythroid surface markers and TED during *ex vivo* culture of human HSPCs. Cells from cocultures of BM CD34^+^ HSPCs with MS-5 stromal cells in medium that did and did not contain Epo were analyzed for surface expression of markers by flow cytometry, and percentages within live cells are shown for days 4, 7, and 10 of culture. **(A)** Representative scatter plots showing CD235a expression in cells cultured with and without Epo. **(B)** Quantification of CD235a expression. **(C)** Representative scatter plots of erythroblast in cultures with and without Epo using a flow cytometry strategy developed by Yan et al., 2021^28^. No erythroblast populations are observed in −Epo cultures. **(D)** Plot of CD49d^+^ cells as a percentage of live cells from cultures in medium with or without Epo. **(E)** Representative scatter plots depicting CD49d^+^ cells within live cells cultured with (top) and without (bottom) Epo. **(F)** Plot of CD117^+^ cells as a percentage of live cells from cultures incubated in complete medium with and without Epo. **(G)** Representative scatter plots depicting CD117^+^ cells within live cells from cultures with (top) or without (bottom) Epo. **(H)** Plot of CD36^+^ cells as a percentage of live cells from cultures incubated in complete medium with and without Epo. **(I)** Representative scatter plots depicting CD36^+^ cells within live cells from cultures with (top) and without (bottom) Epo. P-values ≤ 0.05 were considered significant; *p < 0.05, **p < 0.01, ***p < 0.001 (n = 3 (A-C); n = 5 (D-I)). Abbreviations: ProE; proerythroblast, BasoE; basophilic erythroblast, PolyE; polychromatic erythroblast, OrthoE; orthochromatic erythroblast, Epo; erythropoietin.

**Figure S2.**
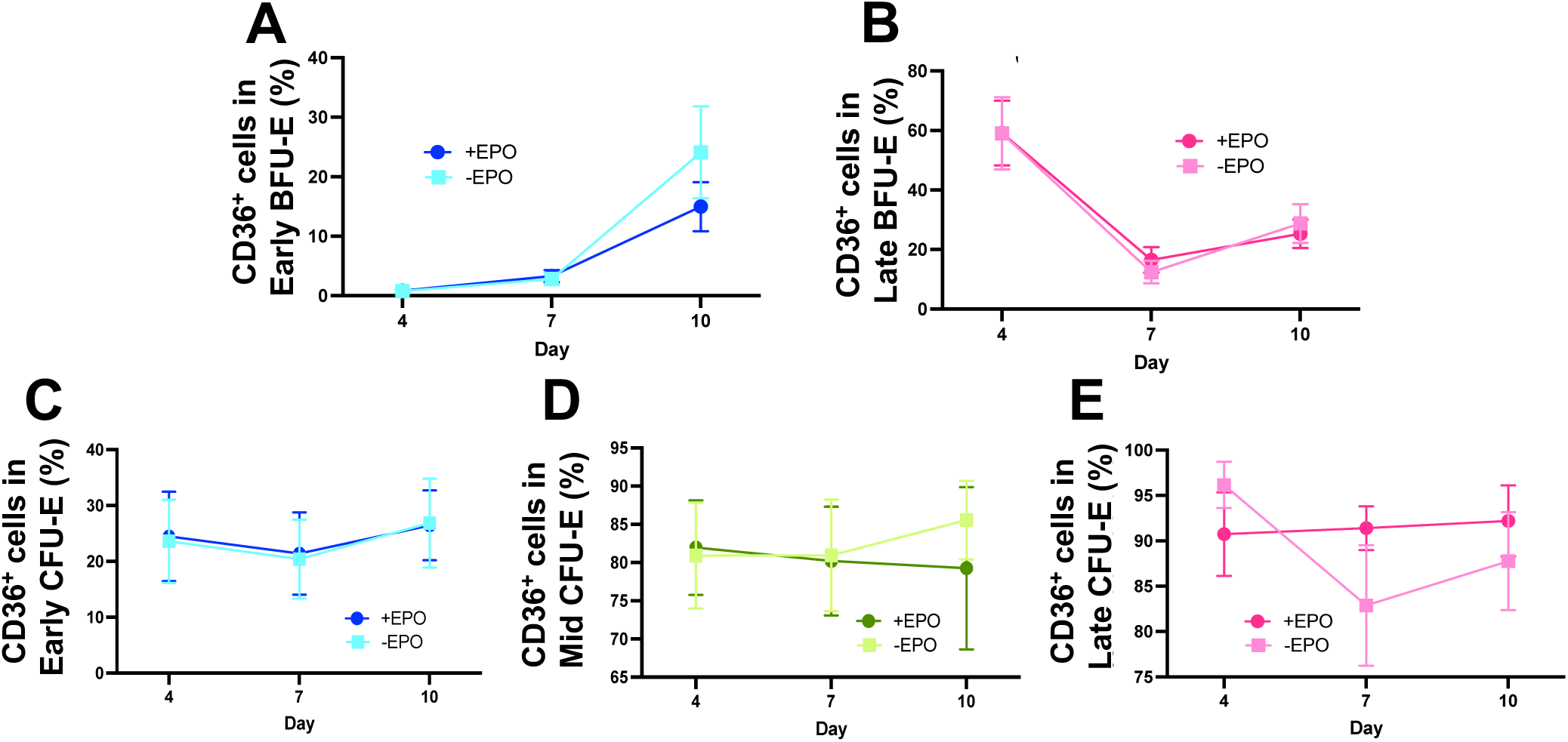
CD36 expression within early erythroid populations. Cells from *ex vivo* cultures of BM CD34^+^ HSPCs incubated in medium containing or lacking Epo were analyzed using the strategy in Figure 2A on days 4, 7, and 10 of culture and CD36 expression was assessed within them (antibodies in Table S1; Panel 1). Line graphs indicate **(H)** CD36^+^ cells within earlyB, **(I)** CD36^+^ cells within lateB, **(J)** CD36^+^ cells within earlyC, **(K)** CD36^+^ cells within midC, and **(K)** CD36^+^ cells within lateC. P-values ≤ 0.05 were considered significant.

**Figure S3.**
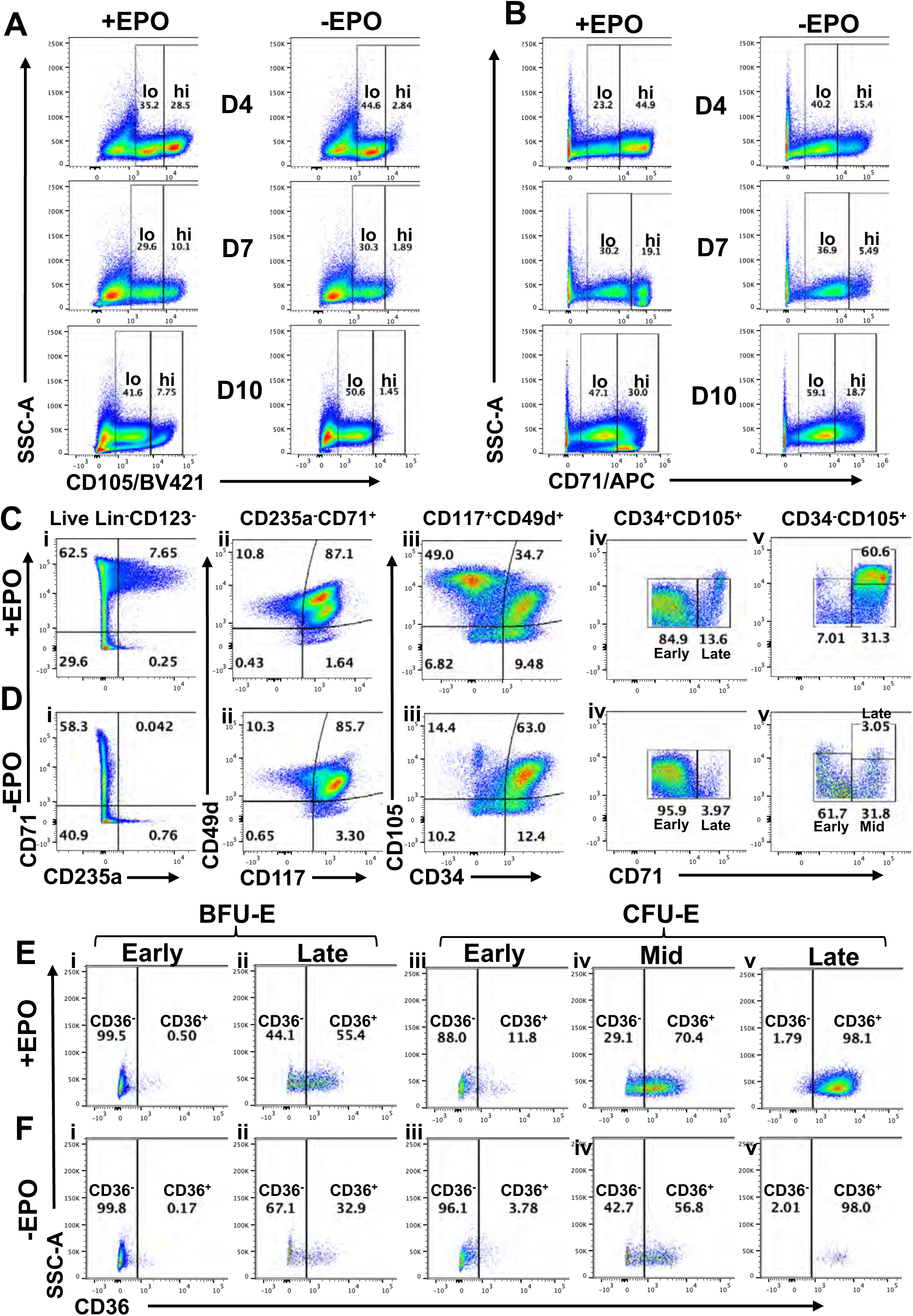
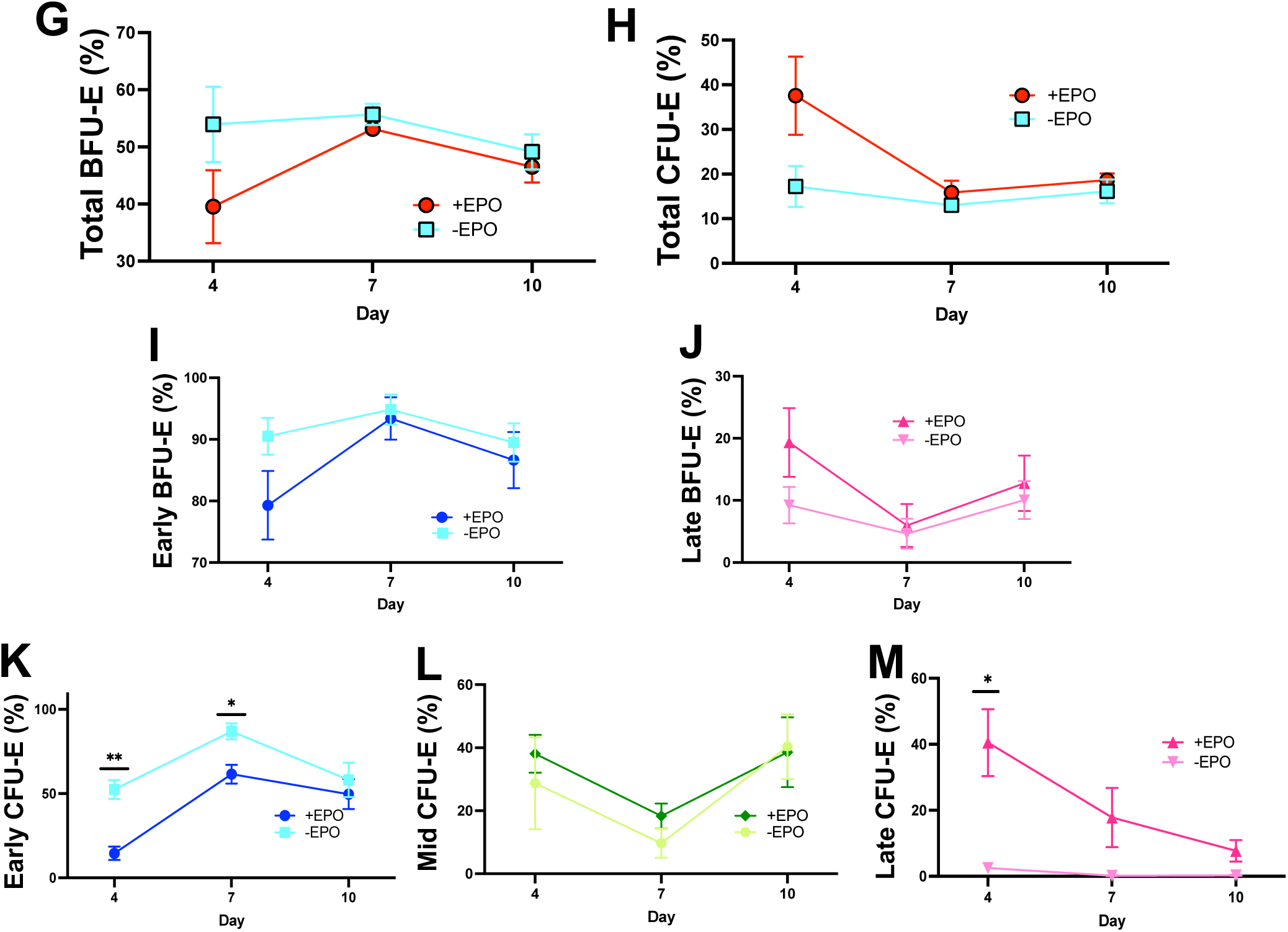
Flow cytometry analysis of heterogeneous BFU-E and CFU-E populations using an alternative antibody panel. Cells from *ex vivo* cultures of BM CD34^+^ HSPCs incubated in medium containing or lacking Epo were analyzed for the indicated surface markers by flow cytometry on days 4, 7, and 10 of culture (antibodies in Table S1; Panel 2). In this analysis, CD105 and CD71 were stained with BV421 and APC labeled antibodies, respectively. The complete list of antibodies is provided in Table S1. **(A)** Representative scatter plots of CD105^+^ cells within live cells. **(B)** Representative scatter plots of CD71^+^ cells within live cells. Cells exhibiting low and high expression are indicated. CD105 and CD71 were stained with BV421 and APC labeled antibodies, respectively. Abbreviation: Epo; erythropoietin. **(C)** Representative scatter plots from the analysis of heterogeneous EE populations in cultures containing Epo. CD235a^+^CD71^+^ cells are gated from LiveLin^-^ cells (i), then CD117^+^CD49d^+^ cells are selected (ii), and CD105^+^ cells are stratified based on CD34 expression (iii). All CD34^+^CD105^+^ cells are considered BFU-Es (iv), and all CD34^−^CD105^+^ cells are considered CFU-Es (v). Further stratification of BFU-E and CFU-E based on CD71 and CD105 expression showed two populations within BFU-Es: CD71^lo^CD105^lo^ (earlyB) and CD71^hi^CD105^lo^ (lateB) (iv). Within CFU-Es, three populations were observed: CD71^lo^CD105^lo^ (earlyC), CD71^hi^CD105^lo^ (midC), and CD71^hi^CD105^hi^ (lateC) (v). **(D)** Representative scatter plots for analysis of populations in cultures lacking Epo by the same strategy as in (A). **(E)** Representative scatter plots depicting CD36 expression within each of the five EE populations from cultures grown in medium containing Epo: i) EarlyB, ii) LateB, iii) EarlyC, iv) MidC, and v) LateC. **(F)** Representative scatter plots depicting CD36 expression within each of the five EE populations from cultures grown in medium lacking Epo: i) EarlyB, ii) LateB, iii) EarlyC, iv) MidC, and v) LateC. Bar graphs indicate percentage of **(G)** Total BFU-E within CD34^+^CD105^+^ cells, **(H)** Total CFU-E within CD34^−^CD105^+^ cells, **(I)** EarlyB within the total BFU-E, **(J)** LateB within the total BFU-E, **(K)** EarlyC within the total CFU-E, **(L)** MidC within the total CFU-E, **(M)** LateC within the total CFU-E. P-values ≤ 0.05 were considered significant (*p < 0.05, **p < 0.01; n = 3).

**Figure S4.**
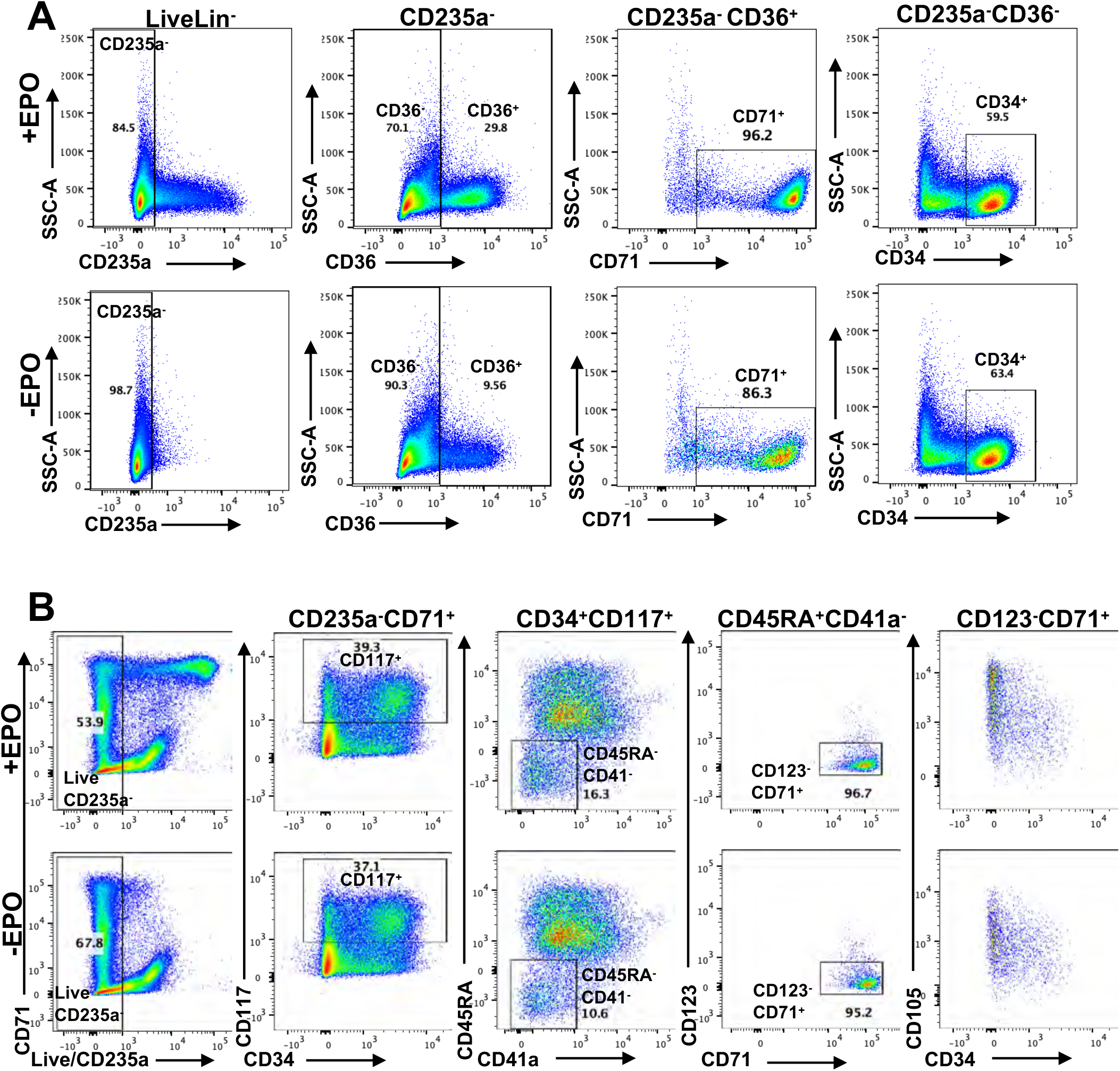
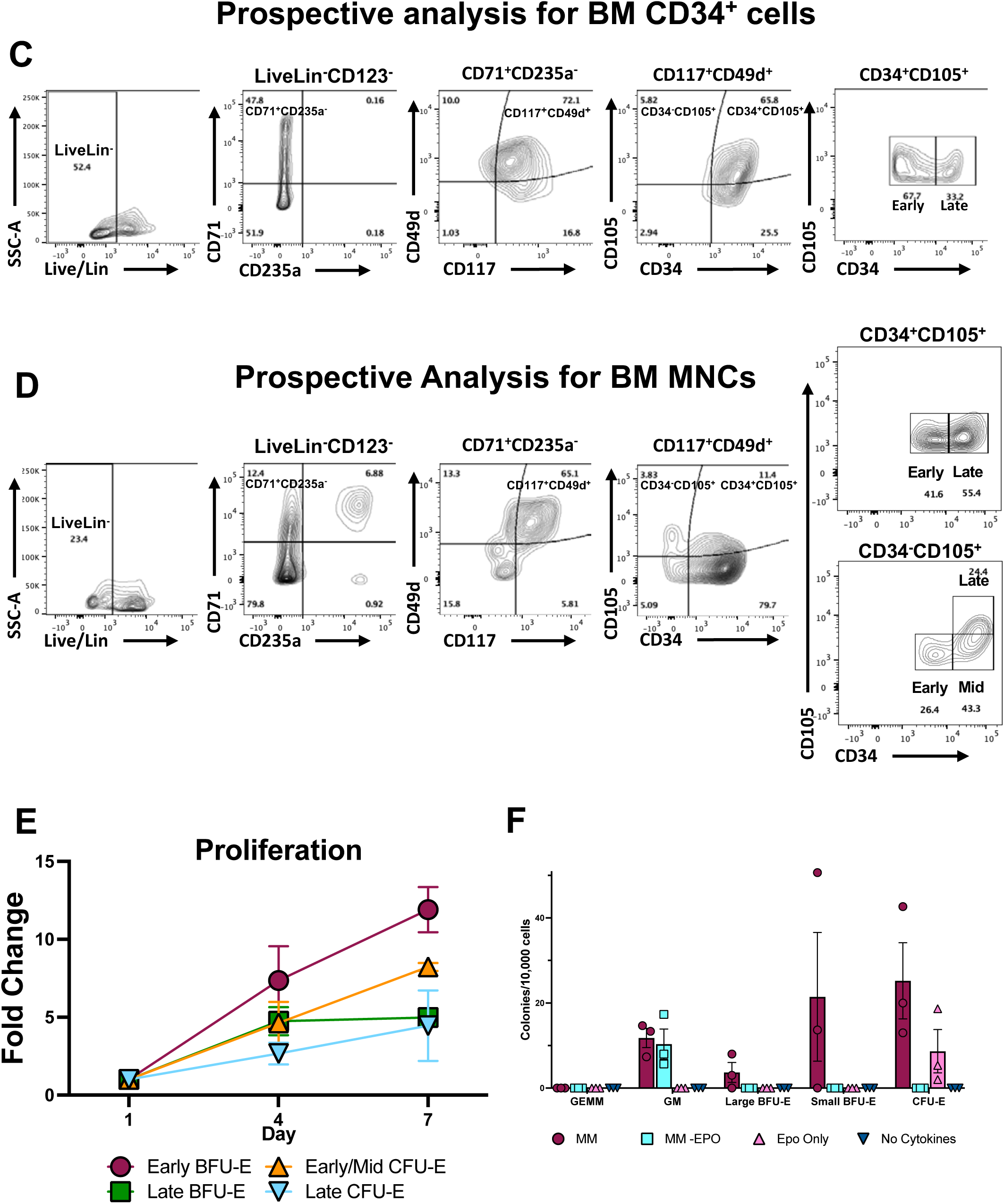
Assessment of early erythroid progenitors using published immunophenotypic strategies and their prospective analysis, and colony forming cell assays. Cells from ex vivo cultures of BM CD34^+^ HSPCs incubated in medium containing or lacking Epo were analyzed for the indicated surface markers by flow cytometry. **(A)** Analysis of EE progenitors using the strategy described by Li et al., 2014^29^. Briefly, CD235a^−^ cells are selected from LiveLin^−^ cells and then stratified based on CD36 expression. CD36^−^ cells were evaluated for CD34 expression, and CD36^−^CD34^+^ cells were considered BFU-Es. CD36^+^ cells are evaluated for CD71 expression, and CD36^−^CD71^+^ cells were considered CFU-Es. **(B)** Assessment of EE progenitors using the immunophenotypic strategy published by Yan et al., 2021^28^. Briefly, CD117^+^ cells were gated from LiveCD235a^−^ cells, then CD45RA^−^CD123^−^ cells were selected and assessed for CD123 and CD71 expression. CD71^+^CD123^−^ cells are then stratified based on expression of CD34 and CD105. All cells in (A-B) were analyzed on day 4 of the *ex vivo* culture. For (C-D), human BM-derived cells were thawed, incubated overnight, and then analyzed using the strategy outlined in Figure 2A. **(C)** Analysis of BM-derived CD34^+^ HSPCs. earlyB and lateB populations were both observed in large numbers. **(D)** Analysis of BM-derived MNCs. All five EE populations were observed. **(E)** Proliferative capacity of sorted early erythroid populations. Cells from *ex vivo* cultures of BM CD34^+^ HSPCs were incubated in a medium containing Epo. On day 4, the five EE subtypes were isolated by FACS and then recultured in suspension. Quantification of cell proliferation for each of the sorted populations (earlyC and midC sorted together) in a complete medium containing Epo is shown (n=4 or 5). **(F)** Colony forming ability of unsorted cells. On day 4, the unsorted cells were plated in methylcellulose containing all cytokines (MethoCult^®^ classic; MM), other cytokine but no Epo (classic without Epo; MM-Epo), Epo-only), or no cytokines. Quantification of colonies (CFU-GEMM, CFU-GM, BFU-E, Large CFU-E, and CFU-E) on day 14 after plating is shown (n = 3). CFU-GM colonies were observed in MM and MM-Epo. Large and small BFU-E colonies were observed only in MM. CFU-E colonies were formed in MM and Epo only medium. No colonies formed in medium lacking cytokines. Abbreviations: EPO; erythropoietin, CFU-GEMM; colony forming unit-granulocyte, erythrocyte, monocyte, megakaryocyte, CFU-GM; colony forming unit-granulocyte monocyte, BFU-E; burst-forming unit erythroid, CFU-E; colony-forming unit erythroid, myeloid media (MM).

**Figure S5.**
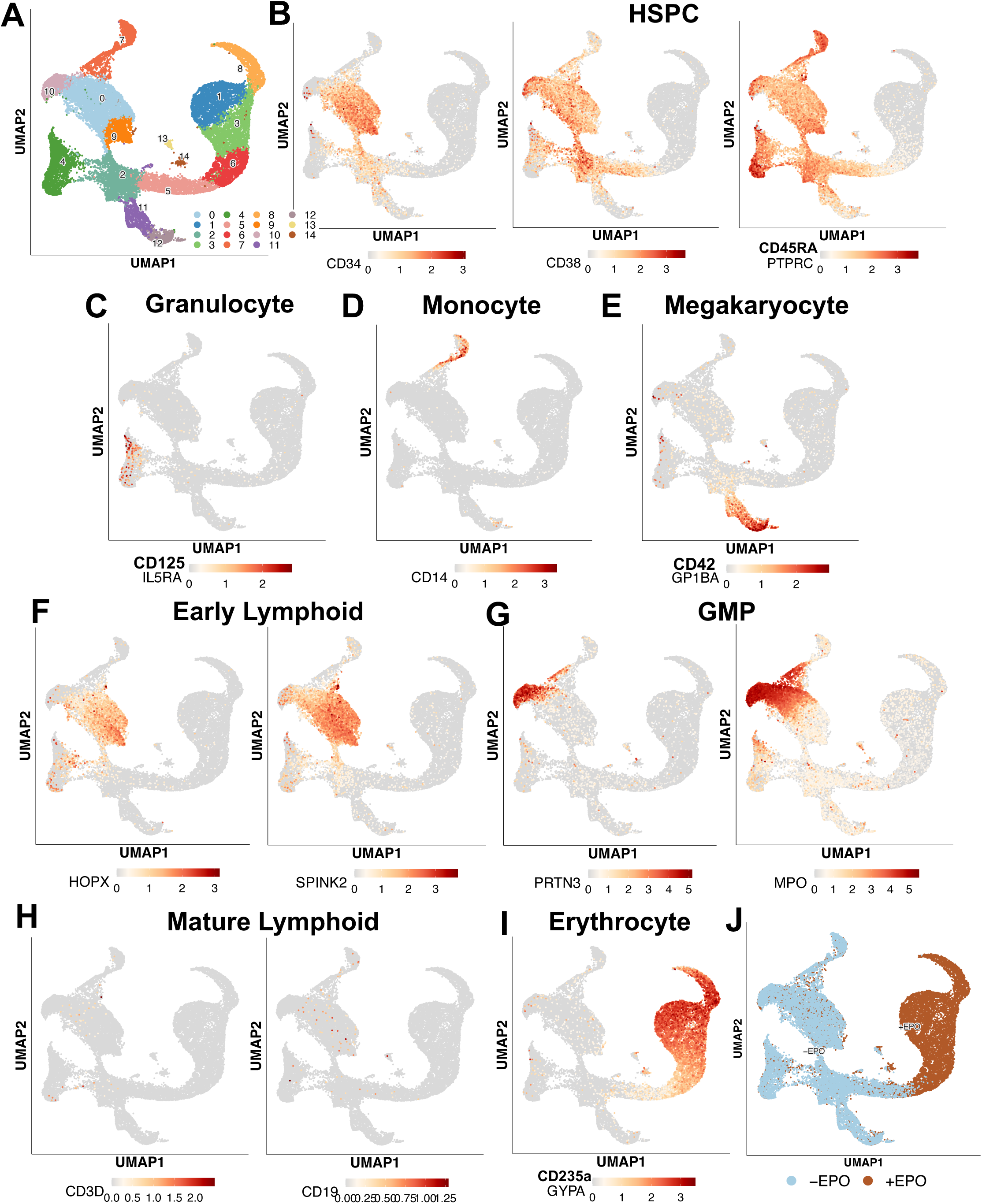
Single cell transcriptomic analysis of hematopoietic markers. UMAP visualization of unsupervised clustering shows cell types (colored by cluster) present in cultures of human BM CD34^+^ HSPCs (**A**). UMAP plots showing expression of markers specific to HSPCs (**B**), granulocytes (**C**), megakaryocytes (**D**), **m**onocytes (**E**), early lymphoid cells (**F**), GMP (**G**), mature lymphoid cells (**H**), and erythrocytes (**I**). UMAP plot indicating cells from cultures grown in the presence and absence of Epo (**J**).

**Figure S6.**
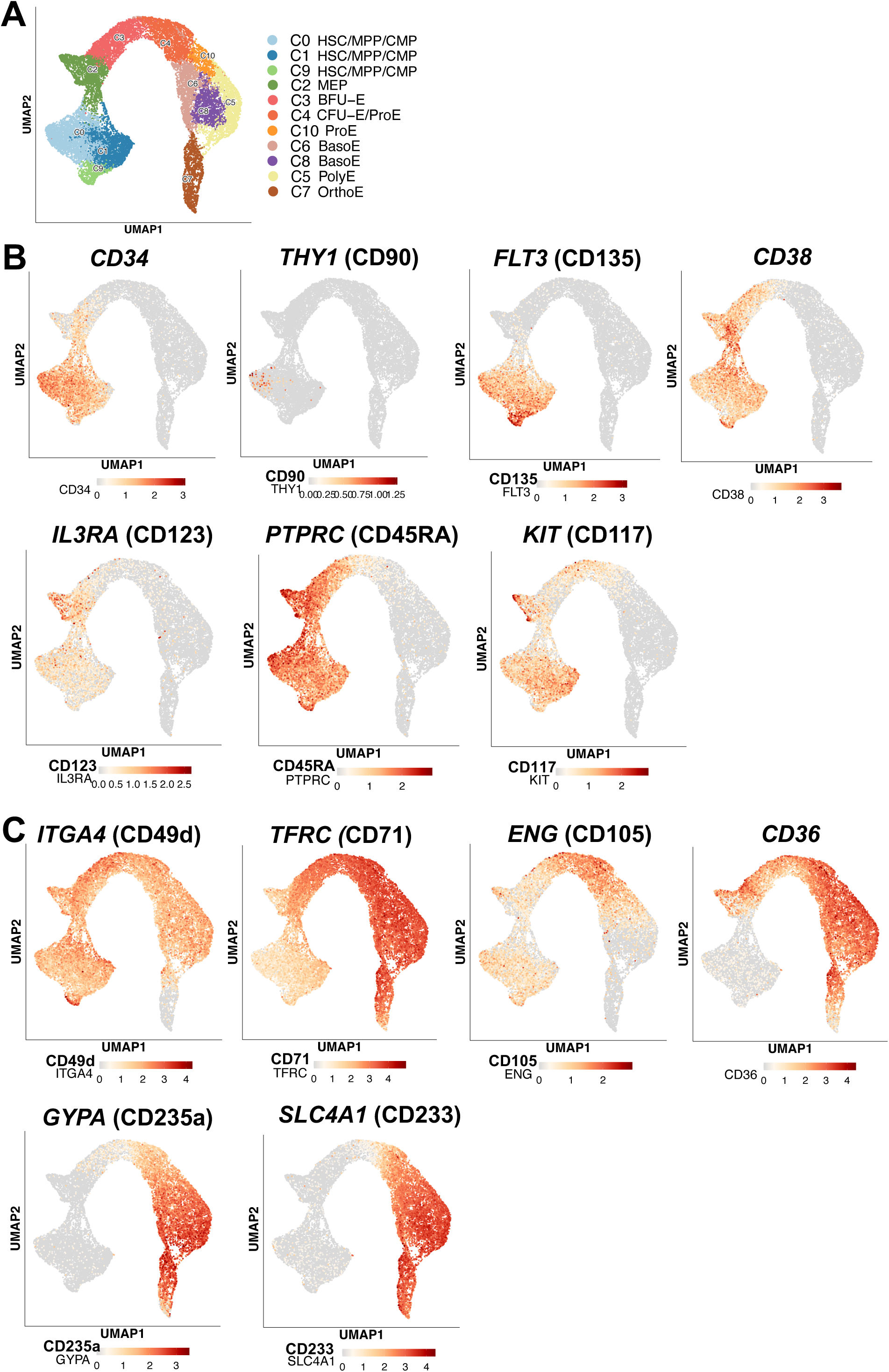
Single cell expression patterns of the HSPC and erythroid genes. UMAP visualization of cells involved in erythropoiesis colored by cluster (**A**), scaled expression of HSPC genes (**B**), and erythroid marker genes (**C**).

**Figure S7.**
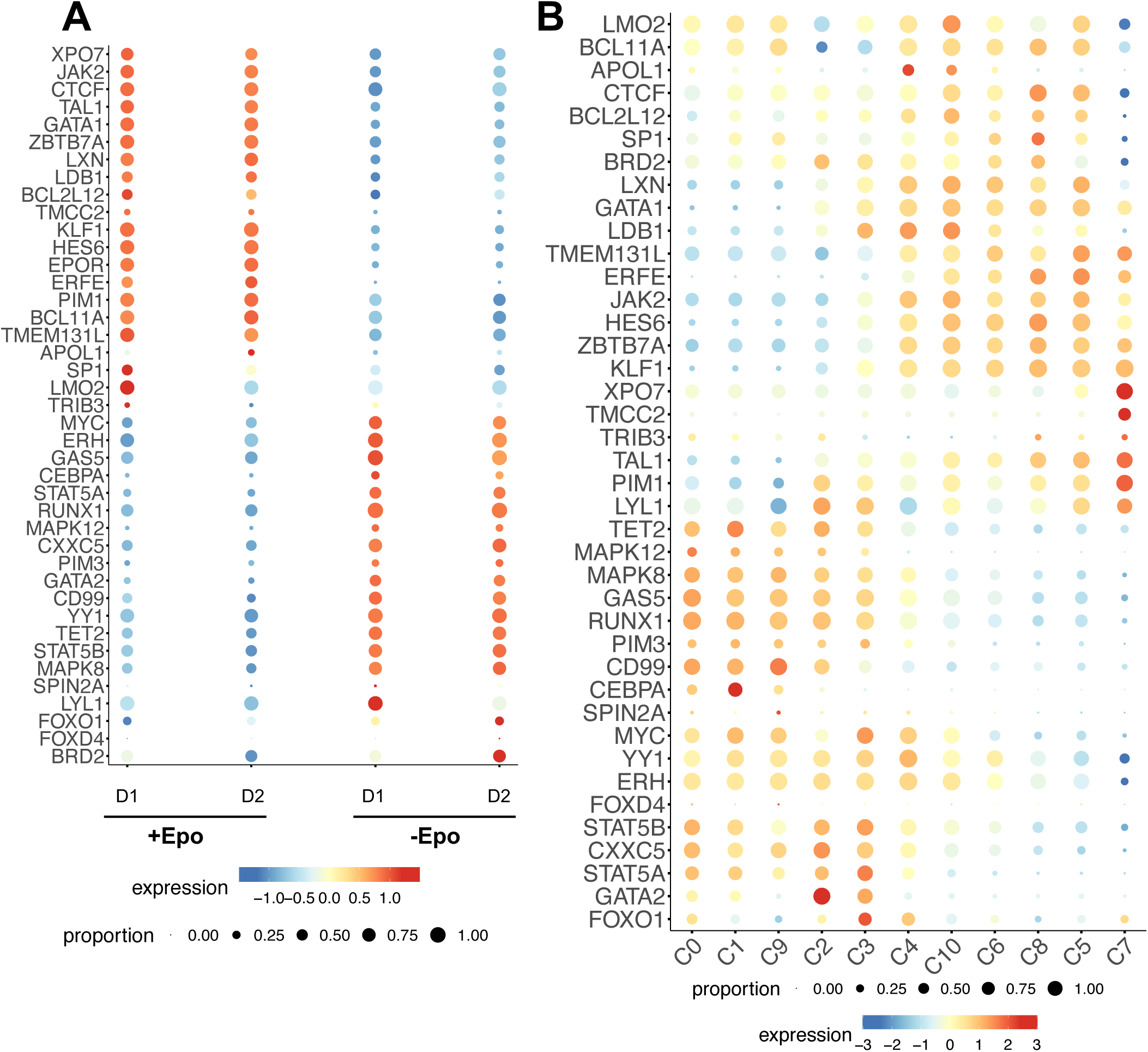
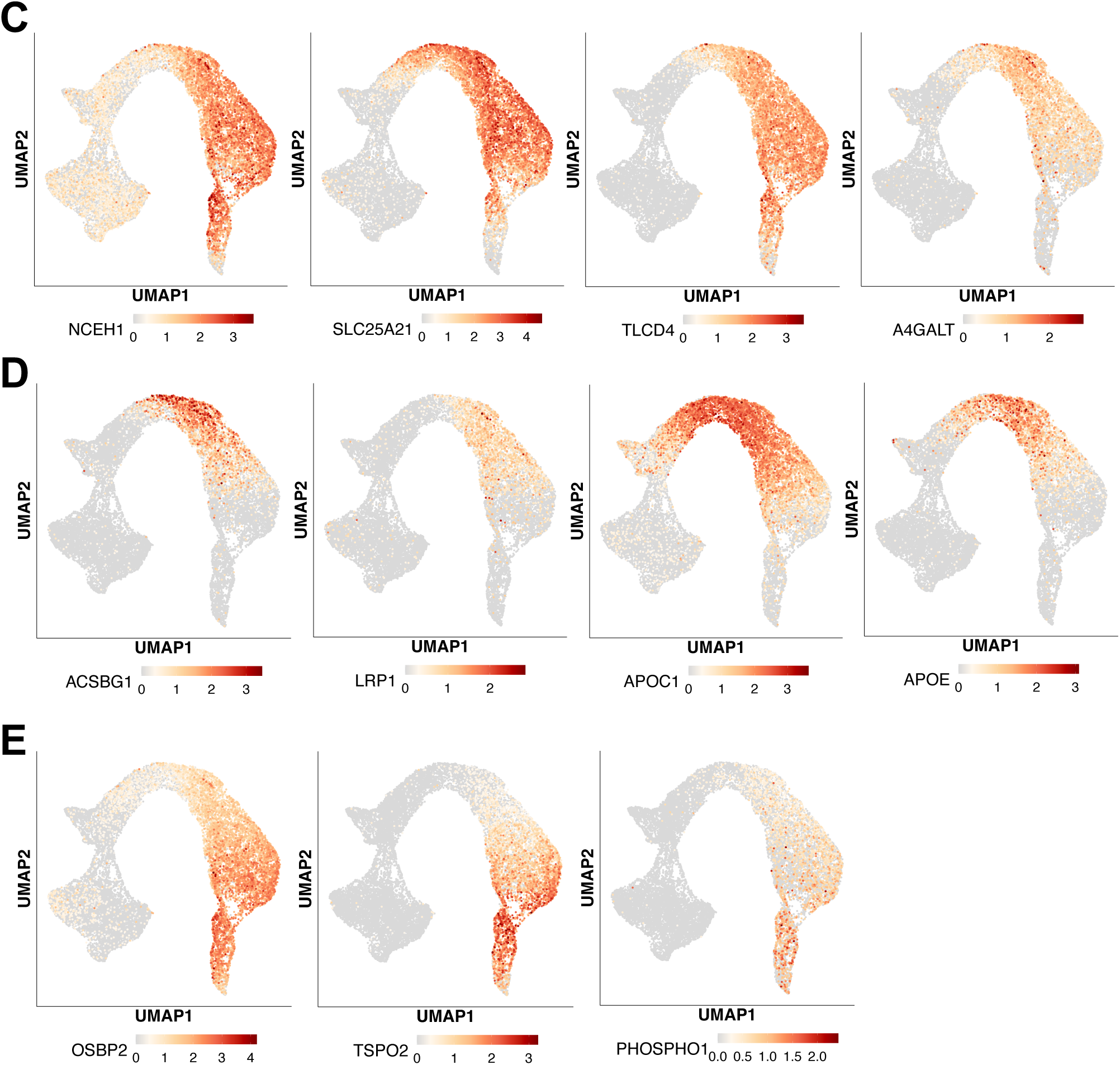
Single cell expression pattern of Epo-dependent genes previously implicated in erythroid differentiation. **(A)** Bubble plot demonstrating the Epo-dependence of genes in cells from both donors. **(B)** Bubble plot showing expression patterns in HSPC and erythroid clusters. Dot color represents normalized expression, and size indicates the proportion of cells that express the gene within a cluster. UMAP plots showing scaled expression of newly identified Epo-dependent lipid metabolism genes that are induced in CFU-Es and expressed through TED (**A**), transiently expressed in EE cells and ProEs (**B**), or later during TED (**C**).

## References

1. Bunn, H.F. (2013). Erythropoietin. Cold Spring Harb Perspect Med 3, a011619. 10.1101/cshperspect.a011619.

2. Bhoopalan, S.V., Huang, L.J., and Weiss, M.J. (2020). Erythropoietin regulation of red blood cell production: from bench to bedside and back. F1000Res *9*. 10.12688/f1000research.26648.1.

3. Valent, P., Busche, G., Theurl, I., Uras, I.Z., Germing, U., Stauder, R., Sotlar, K., Fureder, W., Bettelheim, P., Pfeilstocker, M., et al. (2018). Normal and pathological erythropoiesis in adults: from gene regulation to targeted treatment concepts. Haematologica 103, 1593–1603. 10.3324/haematol.2018.192518.

4. Grover, A., Mancini, E., Moore, S., Mead, A.J., Atkinson, D., Rasmussen, K.D., O’Carroll, D., Jacobsen, S.E., and Nerlov, C. (2014). Erythropoietin guides multipotent hematopoietic progenitor cells toward an erythroid fate. J Exp Med 211, 181–188. 10.1084/jem.20131189.

5. Iscove, N.N., and Guilbert, L.J. (1978). Erythropoietin-Independence of Early Erythropoiesis and a Two-Regulator Model of Proliferative Control in the Hemopoietic System. In In Vitro Aspects of Erythropoiesis, M.J. Murphy, C. Peschle, A.S. Gordon, and E.A. Mirand, eds. (Springer US), pp. 3-7. 10.1007/978-1-4612-6301-2_2.

6. Lin, C.S., Lim, S.K., D’Agati, V., and Costantini, F. (1996). Differential effects of an erythropoietin receptor gene disruption on primitive and definitive erythropoiesis. Genes Dev 10, 154–164. 10.1101/gad.10.2.154.

7. Malik, J., Kim, A.R., Tyre, K.A., Cherukuri, A.R., and Palis, J. (2013). Erythropoietin critically regulates the terminal maturation of murine and human primitive erythroblasts. Haematologica 98, 1778–1787. 10.3324/haematol.2013.087361.

8. Kieran, M.W., Perkins, A.C., Orkin, S.H., and Zon, L.I. (1996). Thrombopoietin rescues in vitro erythroid colony formation from mouse embryos lacking the erythropoietin receptor. Proc Natl Acad Sci U S A 93, 9126–9131. 10.1073/pnas.93.17.9126.

9. Wu, H., Liu, X., Jaenisch, R., and Lodish, H.F. (1995). Generation of committed erythroid BFU-E and CFU-E progenitors does not require erythropoietin or the erythropoietin receptor. Cell 83, 59–67. 10.1016/0092-8674(95)90234-1.

10. Mulcahy, L. (2001). The erythropoietin receptor. Semin Oncol 28, 19–23. 10.1016/s0093-7754(01)90208-8.

11. Witthuhn, B.A., Quelle, F.W., Silvennoinen, O., Yi, T., Tang, B., Miura, O., and Ihle, J.N. (1993). JAK2 associates with the erythropoietin receptor and is tyrosine phosphorylated and activated following stimulation with erythropoietin. Cell 74, 227–236. 10.1016/0092-8674(93)90414-l.

12. Tsiftsoglou, A.S. (2021). Erythropoietin (EPO) as a Key Regulator of Erythropoiesis, Bone Remodeling and Endothelial Transdifferentiation of Multipotent Mesenchymal Stem Cells (MSCs): Implications in Regenerative Medicine. Cells 10. 10.3390/cells10082140.

13. Tothova, Z., Semelakova, M., Solarova, Z., Tomc, J., Debeljak, N., and Solar, P. (2021). The Role of PI3K/AKT and MAPK Signaling Pathways in Erythropoietin Signalization. Int J Mol Sci 22. 10.3390/ijms22147682.

14. Tothova, Z., Tomc, J., Debeljak, N., and Solar, P. (2021). STAT5 as a Key Protein of Erythropoietin Signalization. Int J Mol Sci 22. 10.3390/ijms22137109.

15. Kuhrt, D., and Wojchowski, D.M. (2015). Emerging EPO and EPO receptor regulators and signal transducers. Blood 125, 3536–3541. 10.1182/blood-2014-11-575357.

16. Schippel, N., and Sharma, S. (2023). Dynamics of human hematopoietic stem and progenitor cell differentiation to the erythroid lineage. Exp Hematol 123, 1–17. 10.1016/j.exphem.2023.05.001.

17. Tur, S., Palii, C.G., and Brand, M. (2024). Cell fate decision in erythropoiesis: Insights from multiomics studies. Exp Hematol 131, 104167. 10.1016/j.exphem.2024.104167.

18. Hattangadi, S.M., Wong, P., Zhang, L., Flygare, J., and Lodish, H.F. (2011). From stem cell to red cell: regulation of erythropoiesis at multiple levels by multiple proteins, RNAs, and chromatin modifications. Blood 118, 6258–6268. 10.1182/blood-2011-07-356006.

19. Dzierzak, E., and Philipsen, S. (2013). Erythropoiesis: development and differentiation. Cold Spring Harb Perspect Med 3, a011601. 10.1101/cshperspect.a011601.

20. An, X., Schulz, V.P., Mohandas, N., and Gallagher, P.G. (2015). Human and murine erythropoiesis. Curr Opin Hematol 22, 206–211. 10.1097/MOH.0000000000000134.

21. Velten, L., Haas, S.F., Raffel, S., Blaszkiewicz, S., Islam, S., Hennig, B.P., Hirche, C., Lutz, C., Buss, E.C., Nowak, D., et al. (2017). Human haematopoietic stem cell lineage commitment is a continuous process. Nat Cell Biol 19, 271–281. 10.1038/ncb3493.

22. Notta, F., Zandi, S., Takayama, N., Dobson, S., Gan, O.I., Wilson, G., Kaufmann, K.B., McLeod, J., Laurenti, E., Dunant, C.F., et al. (2016). Distinct routes of lineage development reshape the human blood hierarchy across ontogeny. Science 351, aab2116. 10.1126/science.aab2116.

23. Belluschi, S., Calderbank, E.F., Ciaurro, V., Pijuan-Sala, B., Santoro, A., Mende, N., Diamanti, E., Sham, K.Y.C., Wang, X., Lau, W.W.Y., et al. (2018). Myelo-lymphoid lineage restriction occurs in the human haematopoietic stem cell compartment before lymphoid-primed multipotent progenitors. Nat Commun 9, 4100. 10.1038/s41467-018-06442-4.

24. Zeng, A.G.X., Iacobucci, I., Shah, S., Mitchell, A., Wong, G., Bansal, S., Gao, Q., Kim, H., Kennedy, J.A., Minden, M.D., et al. (2023). Precise single-cell transcriptomic mapping of normal and leukemic cell states reveals unconventional lineage priming in acute myeloid leukemia. bioRxiv. 10.1101/2023.12.26.573390.

25. Ainciburu, M., Ezponda, T., Berastegui, N., Alfonso-Pierola, A., Vilas-Zornoza, A., San Martin-Uriz, P., Alignani, D., Lamo-Espinosa, J., San-Julian, M., Jimenez-Solas, T., et al. (2023). Uncovering perturbations in human hematopoiesis associated with healthy aging and myeloid malignancies at single-cell resolution. Elife 12. 10.7554/eLife.79363.

26. Palii, C.G., Cheng, Q., Gillespie, M.A., Shannon, P., Mazurczyk, M., Napolitani, G., Price, N.D., Ranish, J.A., Morrissey, E., Higgs, D.R., and Brand, M. (2019). Single-Cell Proteomics Reveal that Quantitative Changes in Co-expressed Lineage-Specific Transcription Factors Determine Cell Fate. Cell Stem Cell 24, 812–820 e815. 10.1016/j.stem.2019.02.006.

27. Yan, H., Hale, J., Jaffray, J., Li, J., Wang, Y., Huang, Y., An, X., Hillyer, C., Wang, N., Kinet, S., et al. (2018). Developmental differences between neonatal and adult human erythropoiesis. Am J Hematol 93, 494–503. 10.1002/ajh.25015.

28. Yan, H., Ali, A., Blanc, L., Narla, A., Lane, J.M., Gao, E., Papoin, J., Hale, J., Hillyer, C.D., Taylor, N., et al. (2021). Comprehensive phenotyping of erythropoiesis in human bone marrow: Evaluation of normal and ineffective erythropoiesis. Am J Hematol 96, 1064–1076. 10.1002/ajh.26247.

29. Li, J., Hale, J., Bhagia, P., Xue, F., Chen, L., Jaffray, J., Yan, H., Lane, J., Gallagher, P.G., Mohandas, N., et al. (2014). Isolation and transcriptome analyses of human erythroid progenitors: BFU-E and CFU-E. Blood 124, 3636–3645. 10.1182/blood-2014-07-588806.

30. Hu, J., Liu, J., Xue, F., Halverson, G., Reid, M., Guo, A., Chen, L., Raza, A., Galili, N., Jaffray, J., et al. (2013). Isolation and functional characterization of human erythroblasts at distinct stages: implications for understanding of normal and disordered erythropoiesis in vivo. Blood 121, 3246–3253. 10.1182/blood-2013-01-476390.

31. Wangen, J.R., Eidenschink Brodersen, L., Stolk, T.T., Wells, D.A., and Loken, M.R. (2014). Assessment of normal erythropoiesis by flow cytometry: important considerations for specimen preparation. Int J Lab Hematol 36, 184–196. 10.1111/ijlh.12151.

32. Mori, Y., Chen, J.Y., Pluvinage, J.V., Seita, J., and Weissman, I.L. (2015). Prospective isolation of human erythroid lineage-committed progenitors. Proc Natl Acad Sci U S A 112, 9638–9643. 10.1073/pnas.1512076112.

33. Machherndl-Spandl, S., Suessner, S., Danzer, M., Proell, J., Gabriel, C., Lauf, J., Sylie, R., Klein, H.U., Bene, M.C., Weltermann, A., and Bettelheim, P. (2013). Molecular pathways of early CD105-positive erythroid cells as compared with CD34-positive common precursor cells by flow cytometric cell-sorting and gene expression profiling. Blood Cancer J 3, e100. 10.1038/bcj.2012.45.

34. Loken, M.R., Shah, V.O., Dattilio, K.L., and Civin, C.I. (1987). Flow cytometric analysis of human bone marrow: I. Normal erythroid development. Blood 69, 255–263.

35. Fajtova, M., Kovarikova, A., Svec, P., Kankuri, E., and Sedlak, J. (2013). Immunophenotypic profile of nucleated erythroid progenitors during maturation in regenerating bone marrow. Leuk Lymphoma 54, 2523–2530. 10.3109/10428194.2013.781167.

36. Iskander, D., Psaila, B., Gerrard, G., Chaidos, A., En Foong, H., Harrington, Y., Karnik, L.C., Roberts, I., de la Fuente, J., and Karadimitris, A. (2015). Elucidation of the EP defect in Diamond-Blackfan anemia by characterization and prospective isolation of human EPs. Blood 125, 2553–2557. 10.1182/blood-2014-10-608042.

37. Bapat, A., Schippel, N., Shi, X., Jasbi, P., Gu, H., Kala, M., Sertil, A., and Sharma, S. (2021). Hypoxia promotes erythroid differentiation through the development of progenitors and proerythroblasts. Exp Hematol 97, 32–46 e35. 10.1016/j.exphem.2021.02.012.

38. Bapat, A., Keita, N., and Sharma, S. (2019). Pan-myeloid Differentiation of Human Cord Blood Derived CD34+ Hematopoietic Stem and Progenitor Cells. J Vis Exp. 10.3791/59836.

39. Bapat, A., Keita, N., Martelly, W., Kang, P., Seet, C., Jacobsen, J.R., Stoilov, P., Hu, C., Crooks, G.M., and Sharma, S. (2018). Myeloid Disease Mutations of Splicing Factor SRSF2 Cause G2-M Arrest and Skewed Differentiation of Human Hematopoietic Stem and Progenitor Cells. Stem Cells 36, 1663–1675. 10.1002/stem.2885.

40. Violidaki, D., Axler, O., Jafari, K., Bild, F., Nilsson, L., Mazur, J., Ehinger, M., and Porwit, A. (2020). Analysis of erythroid maturation in the nonlysed bone marrow with help of radar plots facilitates detection of flow cytometric aberrations in myelodysplastic syndromes. Cytometry B Clin Cytom 98, 399–411. 10.1002/cyto.b.21931.

41. Gautier, E.F., Ducamp, S., Leduc, M., Salnot, V., Guillonneau, F., Dussiot, M., Hale, J., Giarratana, M.C., Raimbault, A., Douay, L., et al. (2016). Comprehensive Proteomic Analysis of Human Erythropoiesis. Cell Rep 16, 1470–1484. 10.1016/j.celrep.2016.06.085.

42. Nakahata, T., and Okumura, N. (1994). Cell surface antigen expression in human erythroid progenitors: erythroid and megakaryocytic markers. Leuk Lymphoma 13, 401–409. 10.3109/10428199409049629.

43. Okumura, N., Tsuji, K., and Nakahata, T. (1992). Changes in cell surface antigen expressions during proliferation and differentiation of human erythroid progenitors. Blood 80, 642–650.

44. Palis, J., and Koniski, A. (2018). Functional Analysis of Erythroid Progenitors by Colony-Forming Assays. Methods Mol Biol 1698, 117–132. 10.1007/978-1-4939-7428-3_7.

45. Gregory, C.J., and Eaves, A.C. (1977). Human marrow cells capable of erythropoietic differentiation in vitro: definition of three erythroid colony responses. Blood 49, 855–864.

46. Ryu, Y., Han, G.H., Jung, E., and Hwang, D. (2023). Integration of Single-Cell RNA-Seq Datasets: A Review of Computational Methods. Mol Cells 46, 106–119. 10.14348/molcells.2023.0009.

47. Mimitou, E.P., Cheng, A., Montalbano, A., Hao, S., Stoeckius, M., Legut, M., Roush, T., Herrera, A., Papalexi, E., Ouyang, Z., et al. (2019). Multiplexed detection of proteins, transcriptomes, clonotypes and CRISPR perturbations in single cells. Nat Methods 16, 409–412. 10.1038/s41592-019-0392-0.

48. Butler, A., Hoffman, P., Smibert, P., Papalexi, E., and Satija, R. (2018). Integrating single-cell transcriptomic data across different conditions, technologies, and species. Nat Biotechnol 36, 411–420. 10.1038/nbt.4096.

49. Zhang, W., Yan, C., Liu, X., Yang, P., Wang, J., Chen, Y., Liu, W., Li, S., Zhang, X., Dong, G., et al. (2022). Global characterization of megakaryocytes in bone marrow, peripheral blood, and cord blood by single-cell RNA sequencing. Cancer Gene Ther 29, 1636–1647. 10.1038/s41417-022-00476-z.

50. Friedman, A.D. (2002). Transcriptional regulation of granulocyte and monocyte development. Oncogene 21, 3377–3390. 10.1038/sj.onc.1205324.

51. Papalexi, E., and Satija, R. (2018). Single-cell RNA sequencing to explore immune cell heterogeneity. Nat Rev Immunol 18, 35–45. 10.1038/nri.2017.76.

52. Huang, P., Zhao, Y., Zhong, J., Zhang, X., Liu, Q., Qiu, X., Chen, S., Yan, H., Hillyer, C., Mohandas, N., et al. (2020). Putative regulators for the continuum of erythroid differentiation revealed by single-cell transcriptome of human BM and UCB cells. Proc Natl Acad Sci U S A 117, 12868–12876. 10.1073/pnas.1915085117.

53. An, X., Schulz, V.P., Li, J., Wu, K., Liu, J., Xue, F., Hu, J., Mohandas, N., and Gallagher, P.G. (2014). Global transcriptome analyses of human and murine terminal erythroid differentiation. Blood 123, 3466–3477. 10.1182/blood-2014-01-548305.

54. Pellin, D., Loperfido, M., Baricordi, C., Wolock, S.L., Montepeloso, A., Weinberg, O.K., Biffi, A., Klein, A.M., and Biasco, L. (2019). A comprehensive single cell transcriptional landscape of human hematopoietic progenitors. Nat Commun 10, 2395. 10.1038/s41467-019-10291-0.

55. Psaila, B., Barkas, N., Iskander, D., Roy, A., Anderson, S., Ashley, N., Caputo, V.S., Lichtenberg, J., Loaiza, S., Bodine, D.M., et al. (2016). Single-cell profiling of human megakaryocyte-erythroid progenitors identifies distinct megakaryocyte and erythroid differentiation pathways. Genome Biol 17, 83. 10.1186/s13059-016-0939-7.

56. Manz, M.G., Miyamoto, T., Akashi, K., and Weissman, I.L. (2002). Prospective isolation of human clonogenic common myeloid progenitors. Proc Natl Acad Sci U S A 99, 11872–11877. 10.1073/pnas.172384399.

57. Subramanian, A., Tamayo, P., Mootha, V.K., Mukherjee, S., Ebert, B.L., Gillette, M.A., Paulovich, A., Pomeroy, S.L., Golub, T.R., Lander, E.S., and Mesirov, J.P. (2005). Gene set enrichment analysis: a knowledge-based approach for interpreting genome-wide expression profiles. Proc Natl Acad Sci U S A 102, 15545–15550. 10.1073/pnas.0506580102.

58. Zhang, Y., Xu, Y., Zhang, S., Lu, Z., Li, Y., and Zhao, B. (2022). The regulation roles of Ca(2+) in erythropoiesis: What have we learned? Exp Hematol 106, 19–30. 10.1016/j.exphem.2021.12.192.

59. Eshghi, S., Vogelezang, M.G., Hynes, R.O., Griffith, L.G., and Lodish, H.F. (2007). Alpha4beta1 integrin and erythropoietin mediate temporally distinct steps in erythropoiesis: integrins in red cell development. J Cell Biol 177, 871–880. 10.1083/jcb.200702080.

60. Nemkov, T., Kingsley, P.D., Dzieciatkowska, M., Malik, J., McGrath, K.E., Hansen, K.C., D’Alessandro, A., and Palis, J. (2022). Circulating primitive murine erythroblasts undergo complex proteomic and metabolomic changes during terminal maturation. Blood Adv 6, 3072–3089. 10.1182/bloodadvances.2021005975.

61. Doty, R.T., Lausted, C.G., Munday, A.D., Yang, Z., Yan, X., Meng, C., Tian, Q., and Abkowitz, J.L. (2023). The transcriptomic landscape of normal and ineffective erythropoiesis at single-cell resolution. Blood Adv 7, 4848–4868. 10.1182/bloodadvances.2023010382.

62. Han, Y., Wang, S., Wang, Y., Huang, Y., Gao, C., Guo, X., Chen, L., Zhao, H., and An, X. (2023). Comprehensive Characterization and Global Transcriptome Analysis of Human Fetal Liver Terminal Erythropoiesis. Genomics Proteomics Bioinformatics 21, 1117–1132. 10.1016/j.gpb.2023.07.001.

63. Kiatpakdee, B., Sato, K., Otsuka, Y., Arashiki, N., Chen, Y., Tsumita, T., Otsu, W., Yamamoto, A., Kawata, R., Yamazaki, J., et al. (2020). Cholesterol-binding protein TSPO2 coordinates maturation and proliferation of terminally differentiating erythroblasts. J Biol Chem 295, 8048–8063. 10.1074/jbc.RA119.011679.

64. Huang, N.J., Lin, Y.C., Lin, C.Y., Pishesha, N., Lewis, C.A., Freinkman, E., Farquharson, C., Millan, J.L., and Lodish, H. (2018). Enhanced phosphocholine metabolism is essential for terminal erythropoiesis. Blood 131, 2955–2966. 10.1182/blood-2018-03-838516.

65. Moura, I.C., Hermine, O., Lacombe, C., and Mayeux, P. (2015). Erythropoiesis and transferrin receptors. Curr Opin Hematol 22, 193–198. 10.1097/MOH.0000000000000133.

66. Richard, C., and Verdier, F. (2020). Transferrin Receptors in Erythropoiesis. Int J Mol Sci 21. 10.3390/ijms21249713.

67. Zhang, L., Magli, A., Catanese, J., Xu, Z., Kyba, M., and Perlingeiro, R.C. (2011). Modulation of TGF-beta signaling by endoglin in murine hemangioblast development and primitive hematopoiesis. Blood 118, 88–97. 10.1182/blood-2010-12-325019.

68. Meurer, S.K., and Weiskirchen, R. (2020). Endoglin: An ‘Accessory’ Receptor Regulating Blood Cell Development and Inflammation. Int J Mol Sci 21. 10.3390/ijms21239247.

69. Schoonderwoerd, M.J.A., Goumans, M.T.H., and Hawinkels, L. (2020). Endoglin: Beyond the Endothelium. Biomolecules 10. 10.3390/biom10020289.

70. Moody, J.L., Singbrant, S., Karlsson, G., Blank, U., Aspling, M., Flygare, J., Bryder, D., and Karlsson, S. (2007). Endoglin is not critical for hematopoietic stem cell engraftment and reconstitution but regulates adult erythroid development. Stem Cells 25, 2809–2819. 10.1634/stemcells.2006-0602.

71. Cho, S.K., Bourdeau, A., Letarte, M., and Zuniga-Pflucker, J.C. (2001). Expression and function of CD105 during the onset of hematopoiesis from Flk1(+) precursors. Blood 98, 3635–3642. 10.1182/blood.v98.13.3635.

72. Buhring, H.J., Muller, C.A., Letarte, M., Gougos, A., Saalmuller, A., van Agthoven, A.J., and Busch, F.W. (1991). Endoglin is expressed on a subpopulation of immature erythroid cells of normal human bone marrow. Leukemia 5, 841–847.

73. Orfao, A., Matarraz, S., Perez-Andres, M., Almeida, J., Teodosio, C., Berkowska, M.A., van Dongen, J.J.M., and EuroFlow (2019). Immunophenotypic dissection of normal hematopoiesis. J Immunol Methods 475, 112684. 10.1016/j.jim.2019.112684.

74. Varricchio, L., Geer, E.B., Martelli, F., Mazzarini, M., Funnell, A., Bieker, J.J., Papayannopoulou, T., and Migliaccio, A.R. (2023). Patients with hypercortisolemic Cushing disease possess a distinct class of hematopoietic progenitor cells leading to erythrocytosis. Haematologica 108, 1053–1067. 10.3324/haematol.2021.280542.

75. Ashley, R.J., Yan, H., Wang, N., Hale, J., Dulmovits, B.M., Papoin, J., Olive, M.E., Udeshi, N.D., Carr, S.A., Vlachos, A., et al. (2020). Steroid resistance in Diamond Blackfan anemia associates with p57Kip2 dysregulation in erythroid progenitors. J Clin Invest 130, 2097–2110. 10.1172/JCI132284.

76. Barrachina, M.N., Pernes, G., Becker, I.C., Allaeys, I., Hirsch, T.I., Groeneveld, D.J., Khan, A.O., Freire, D., Guo, K., Carminita, E., et al. (2023). Efficient megakaryopoiesis and platelet production require phospholipid remodeling and PUFA uptake through CD36. bioRxiv. 10.1101/2023.02.12.527706.

77. Chami, N., Chen, M.H., Slater, A.J., Eicher, J.D., Evangelou, E., Tajuddin, S.M., Love-Gregory, L., Kacprowski, T., Schick, U.M., Nomura, A., et al. (2016). Exome Genotyping Identifies Pleiotropic Variants Associated with Red Blood Cell Traits. Am J Hum Genet 99, 8–21. 10.1016/j.ajhg.2016.05.007.

78. Auer, P.L., Johnsen, J.M., Johnson, A.D., Logsdon, B.A., Lange, L.A., Nalls, M.A., Zhang, G., Franceschini, N., Fox, K., Lange, E.M., et al. (2012). Imputation of exome sequence variants into population-based samples and blood-cell-trait-associated loci in African Americans: NHLBI GO Exome Sequencing Project. Am J Hum Genet 91, 794–808. 10.1016/j.ajhg.2012.08.031.

79. Meng, Y., Pospiech, M., Ali, A., Chandwani, R., Vergel, M., Onyemaechi, S., Yaghmour, G., Lu, R., and Alachkar, H. (2023). Deletion of CD36 exhibits limited impact on normal hematopoiesis and the leukemia microenvironment. Cell Mol Biol Lett 28, 45. 10.1186/s11658-023-00455-8.

80. Alattar, A.G., Storry, J.R., and Olsson, M.L. (2024). Evidence that CD36 is expressed on red blood cells and constitutes a novel blood group system of clinical importance. Vox Sang 119, 496–504. 10.1111/vox.13595.

81. Quintana, A.M., Picchione, F., Klein Geltink, R.I., Taylor, M.R., and Grosveld, G.C. (2014). Zebrafish ETV7 regulates red blood cell development through the cholesterol synthesis pathway. Dis Model Mech 7, 265–270. 10.1242/dmm.012526.

82. Hernandez, J.A., Castro, V.L., Reyes-Nava, N., Montes, L.P., and Quintana, A.M. (2019). Mutations in the zebrafish hmgcs1 gene reveal a novel function for isoprenoids during red blood cell development. Blood Adv 3, 1244–1254. 10.1182/bloodadvances.2018024539.

83. Fan, J., Rone, M.B., and Papadopoulos, V. (2009). Translocator protein 2 is involved in cholesterol redistribution during erythropoiesis. J Biol Chem 284, 30484–30497. 10.1074/jbc.M109.029876.

84. Lu, Z., Huang, L., Li, Y., Xu, Y., Zhang, R., Zhou, Q., Sun, Q., Lu, Y., Chen, J., Shen, Y., et al. (2022). Fine-Tuning of Cholesterol Homeostasis Controls Erythroid Differentiation. Adv Sci (Weinh) 9, e2102669. 10.1002/advs.202102669.

85. Holm, T.M., Braun, A., Trigatti, B.L., Brugnara, C., Sakamoto, M., Krieger, M., and Andrews, N.C. (2002). Failure of red blood cell maturation in mice with defects in the high-density lipoprotein receptor SR-BI. Blood 99, 1817–1824. 10.1182/blood.v99.5.1817.

86. Shalev, H., Kapelushnik, J., Moser, A., Knobler, H., and Tamary, H. (2007). Hypocholesterolemia in chronic anemias with increased erythropoietic activity. Am J Hematol 82, 199–202. 10.1002/ajh.20804.

87. Meurs, I., Hoekstra, M., van Wanrooij, E.J., Hildebrand, R.B., Kuiper, J., Kuipers, F., Hardeman, M.R., Van Berkel, T.J., and Van Eck, M. (2005). HDL cholesterol levels are an important factor for determining the lifespan of erythrocytes. Exp Hematol 33, 1309–1319. 10.1016/j.exphem.2005.07.004.

88. Bernecker, C., Kofeler, H., Pabst, G., Trotzmuller, M., Kolb, D., Strohmayer, K., Trajanoski, S., Holzapfel, G.A., Schlenke, P., and Dorn, I. (2019). Cholesterol Deficiency Causes Impaired Osmotic Stability of Cultured Red Blood Cells. Front Physiol 10, 1529. 10.3389/fphys.2019.01529.

89. Saito, K., van der Garde, M., Umemoto, T., Miharada, N., Sjoberg, J., Sigurdsson, V., Shirozu, H., Kamei, S., Radulovic, V., Suzuki, M., et al. (2024). Lipoprotein metabolism mediates hematopoietic stem cell responses under acute anemic conditions. Nat Commun 15, 8131. 10.1038/s41467-024-52509-w.

90. Weiss, G., Ganz, T., and Goodnough, L.T. (2019). Anemia of inflammation. Blood 133, 40–50. 10.1182/blood-2018-06-856500.

91. Hess, S.Y., Owais, A., Jefferds, M.E.D., Young, M.F., Cahill, A., and Rogers, L.M. (2023). Accelerating action to reduce anemia: Review of causes and risk factors and related data needs. Ann N Y Acad Sci 1523, 11–23. 10.1111/nyas.14985.

92. Lewis, R., Bewersdorf, J.P., and Zeidan, A.M. (2021). Clinical Management of Anemia in Patients with Myelodysplastic Syndromes: An Update on Emerging Therapeutic Options. Cancer Manag Res 13, 645–657. 10.2147/CMAR.S240600.

93. Garcia-Manero, G., Santini, V., Almeida, A., Platzbecker, U., Jonasova, A., Silverman, L.R., Falantes, J., Reda, G., Buccisano, F., Fenaux, P., et al. (2021). Phase III, Randomized, Placebo-Controlled Trial of CC-486 (Oral Azacitidine) in Patients With Lower-Risk Myelodysplastic Syndromes. J Clin Oncol *39*, 1426-1436. 10.1200/JCO.20.02619.

94. Zheng, G.X., Terry, J.M., Belgrader, P., Ryvkin, P., Bent, Z.W., Wilson, R., Ziraldo, S.B., Wheeler, T.D., McDermott, G.P., Zhu, J., et al. (2017). Massively parallel digital transcriptional profiling of single cells. Nat Commun 8, 14049. 10.1038/ncomms14049.

95. Hao, Y., Hao, S., Andersen-Nissen, E., Mauck, W.M., 3rd, Zheng, S., Butler, A., Lee, M.J., Wilk, A.J., Darby, C., Zager, M., et al. (2021). Integrated analysis of multimodal single-cell data. Cell 184, 3573-3587 e3529. 10.1016/j.cell.2021.04.048.

96. McGinnis, C.S., Murrow, L.M., and Gartner, Z.J. (2019). DoubletFinder: Doublet Detection in Single-Cell RNA Sequencing Data Using Artificial Nearest Neighbors. Cell Syst 8, 329–337 e324. 10.1016/j.cels.2019.03.003.

97. Ouyang, J.F., Kamaraj, U.S., Cao, E.Y., and Rackham, O.J.L. (2021). ShinyCell: simple and sharable visualization of single-cell gene expression data. Bioinformatics 37, 3374–3376. 10.1093/bioinformatics/btab209.

98. Trapnell, C., Cacchiarelli, D., Grimsby, J., Pokharel, P., Li, S., Morse, M., Lennon, N.J., Livak, K.J., Mikkelsen, T.S., and Rinn, J.L. (2014). The dynamics and regulators of cell fate decisions are revealed by pseudotemporal ordering of single cells. Nat Biotechnol 32, 381–386. 10.1038/nbt.2859.

99. Cao, J., Spielmann, M., Qiu, X., Huang, X., Ibrahim, D.M., Hill, A.J., Zhang, F., Mundlos, S., Christiansen, L., Steemers, F.J., et al. (2019). The single-cell transcriptional landscape of mammalian organogenesis. Nature 566, 496–502. 10.1038/s41586-019-0969-x.

100. Elizarraras, J.M., Liao, Y., Shi, Z., Zhu, Q., Pico, A.R., and Zhang, B. (2024). WebGestalt 2024: faster gene set analysis and new support for metabolomics and multi-omics. Nucleic Acids Res 52, W415–W421. 10.1093/nar/gkae456.

